# Multiscale characterisation of the human claustrum from histology to MRI

**DOI:** 10.64898/2025.12.06.692728

**Authors:** Navona Calarco, Skerdi Progri, Sriranga Kashyap, Shuting Xie, Claude Lepage, Boris C. Bernhardt, Alan C. Evans, Kâmil Uludağ

## Abstract

Though linked to an unusually broad array of functions, the human claustrum’s complex morphology has hindered *in vivo* study, resulting in a small MRI literature marked by implausibly large discrepancies in reported characteristics. We constructed the first three-dimensional histological “gold standard” claustrum model, and systematically evaluated *in vivo* 7-Tesla MRI datasets against it and downsampled derivatives. MRI showed resolution-dependent differences rather than contrast limitations, transforming the claustrum’s intricate sheet into an artefactually thickened ribbon. However, submillimetre MRI reliably recovered the dorsal “core” that contains most claustral volume and density and houses major corticoclaustral connectivity. At 0.5mm resolution, extension into the temporal lobe, including irregular ventral “puddles”, was partially recovered, with uncertainty reflecting boundary imprecision rather than anatomical loss. Our results refute the view that the claustrum is inaccessible in the living human brain, define practical measurement limits, and provide a foundation for future functional investigations.

## INTRODUCTION

Two decades ago, Crick and Koch argued that the claustrum’s widespread cortical connectivity made it a candidate neural correlate of consciousness, igniting modern claustrum research^1^. Since, animal work has elaborated the claustrum’s extensive connectivity^2^, and inventive experiments implicate it in a diverse array of functions^3^ so expansive that only basic sensory and motor processing remain outside its remit^4^.

Despite this progress, the function of the human claustrum remains elusive, as methodological barriers have long stymied *in vivo* study. Crick and Koch lamented that MRI lacked the resolution needed to capture the claustrum’s irregular geometry. The dorsal claustrum is extremely thin mediolaterally and separated from the putamen and insula by only a slender white-matter band. The ventral claustrum broadens as it nears both the piriform and amygdaloid complex but exhibits lower density with cell dispersion through irregular fibre spaces^5–7^. In principle, these features fall well below the nominal voxel size of structural MRI afforded by conventional and high magnetic field strengths (i.e. 1.5 and 3-Tesla), and perhaps also ultra-high field imaging (>7-Tesla)^8^.

Still, a small human MRI literature has emerged. Limitations are evident: published images show clear partial voluming with adjacent capsules and nearby cortical and subcortical structures. Few studies report quantitative metrics, but 13 providing volume estimates in healthy adults differ by over fourfold, far exceeding typical within-subject variability^9^, and diverging sharply from histology-based estimates (**Fig. 1**). Nonetheless, MRI has yielded insight on the “claustrum sign,” a bilateral hyperintensity on T2-weighted and FLAIR images that is detectable even at low field and coarse resolution, and has long aided diagnosis of Wilson’s disease^10^. Pioneering diffusion and functional MRI studies have extended landmark animal findings suggesting that the claustrum is among the most highly connected structures^11,12^, and may contribute to cognitive control^13–15^, pain perception^16^, and higher-order processing^17^.

**Fig. 1.**
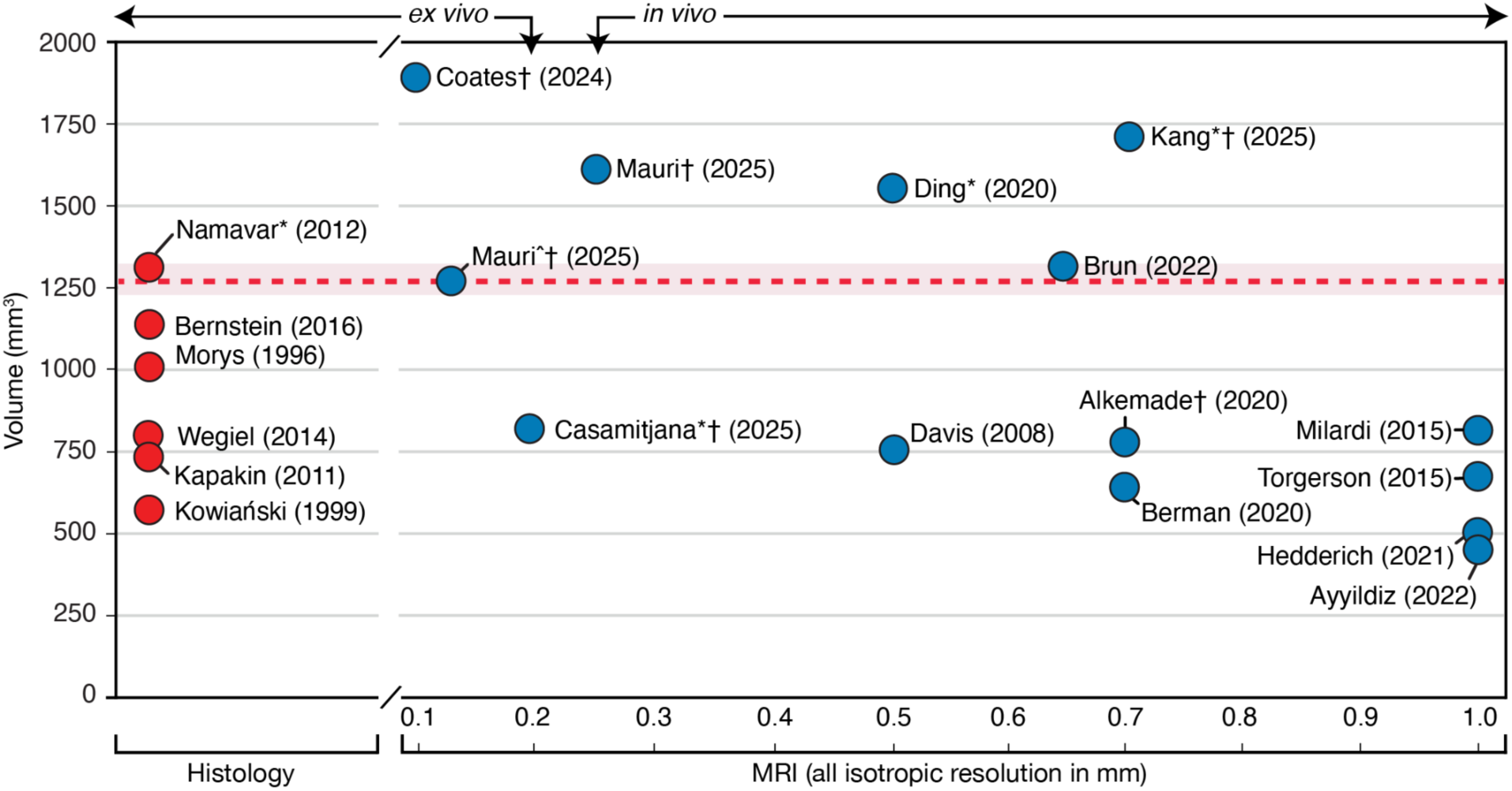
Claustrum volume estimates from past histological and MRI literature. Six histology studies (red) and 13 MRI studies (blue) provided manually or semi-manually segmented claustrum volume estimates for healthy adults. Values were extracted from published tables or figures, available data, or provided by authors upon request, and are shown as reported or available, without harmonisation across methods. The gold-standard estimate is shown as the red dashed line. Across MRI studies, there is more than a four-fold range between the smallest and largest reported volumes, with higher-resolution MRI yielding larger estimates (*r*=–0.62, *p*=0.016). Estimates marked with an asterisk (*) represent a single hemisphere; all others reflect the mean of both hemispheres. Estimates marked with a dagger (†) derive from 7-Tesla MRI. Estimates from Mauri^ (2025) derive from 15 ex vivo scans spanning 0.10–0.25mm isotropic resolution. Where multiple publications analysed the same dataset, the earliest is cited.

Limitations notwithstanding, mapping the claustrum in the living human brain may ultimately rely on MRI. Direct human evidence is exceptionally rare, and animal models face translational barriers; both fail to definitively adjudicate between competing functional hypotheses^4^. Complete bilateral absence is reported in only nine congenital cases, all with widespread atrophy and typically fatal in infancy^18–23^. Acquired lesions are unilateral, incomplete, and/or non-specific^24–27^. Intraoperative stimulation has produced intriguing but inconsistent effects, reflecting opportunistic electrode placement and co-activation of adjacent tissue^28,29^. Rodent models are common but differ markedly from humans: rodents lack an extreme capsule, complicating the insular boundary^30,31^; their endopiriform nucleus is distinct but continuous with the claustrum in humans^30^; the inferior ventral “puddles” prominent in humans are poorly developed^32^, and their claustrum occupies a much larger relative volume^33^. Though rodents exhibit substantial claustral-cortical connectivity^34–36^, functional theories do not cleanly extend to lissencephalic brains with fewer and less differentiated cortical areas.

The critical question is whether MRI can capture this elusive nucleus with the fidelity needed for discovery, beyond coarse disease markers and inference by analogy from animal studies. Ultra-high field scanners are increasingly common (map) and now achieve anatomical isotropic resolutions of 0.7mm, with some implementations reaching 0.5mm^37,38^, enabling fine-grained studies of neocortical networks^39^, and deep structures including the substantia nigra^40^, thalamic subnuclei^41^, auditory nuclei^42^, nucleus basalis^43^, and hippocampus^44,45^. Tissue contrast is unlikely to be limiting: the claustrum is visible on T1-weighted scans despite partial voluming, consistent with its glutamatergic neurons^46^ and low iron and moderate myelin content, which confer cortical-like signal properties^47^. Yet the field remains cautious, with only a handful of claustrum studies acquiring submillimetre voxels, and just two leveraging ultra-high field strength^13,17^.

One factor contributing to the lag in *in vivo* human MRI may be the lack of a high-resolution, three-dimensional histological reference atlas to evaluate MRI’s resolving capacity. Classical anatomical studies are richly descriptive but limited by coronal sectioning with large gaps, challenging imagination of the claustrum’s undulating course^5–7^. One prior study generated a three-dimensional histological model, but it was low resolution, excluded the ventral claustrum, and exists only as photographs^48^. Modern whole-brain digital atlases are more densely sampled, but delineate the claustrum *de novo* without specialist criteria, and diverge radically in their depiction of the ventral extent^49,50^. Two recent studies advanced the field by making publicly available claustrum segmentations from *ex vivo* MRI at 100µm resolution^51,52^, but validating MRI with MRI ultimately begs our present question of if MRI can truly resolve claustral structure.

To advance the broader goal of elucidating human claustral function, we here address the antecedent question of whether MRI can accurately capture this elusive nucleus *in vivo*, using two complementary approaches. First, we segmented the BigBrain dataset^53^ to create the first continuous, high-resolution, histology-based three-dimensional claustrum atlas (a “gold standard”), enabling detailed morphometric description. Second, we compared this atlas and its downsampled derivatives with manual claustrum segmentations from three 7-Tesla datasets (0.5, 0.7, and 1.0 mm isotropic resolution)^38,44,54^. Our approach disentangles spatial sampling effects from other factors, establishes resolution-specific benchmarks, and supplies the missing foundation for next-generation studies of claustral connectivity and function.

## RESULTS

### High-resolution histology reveals an extremely thin and fragmented claustrum

Drawing on the exceptional anatomical detail of the 100µm BigBrain dataset, we manually delineated a continuous bilateral segmentation of the human claustrum surpassing the detail of existing histological atlases (**Fig. 2**). The resulting “gold standard” model recapitulates defining features described in classical literature: dorsally, an exquisitely thin sheet follows the insular convolution and bends laterally over the central insular sulcus; ventrally, the claustrum broadens into a reticular arrangement, fragmenting into small “puddles’’ separated by white-matter laminae in the anterior temporal lobe.

**Fig. 2.**
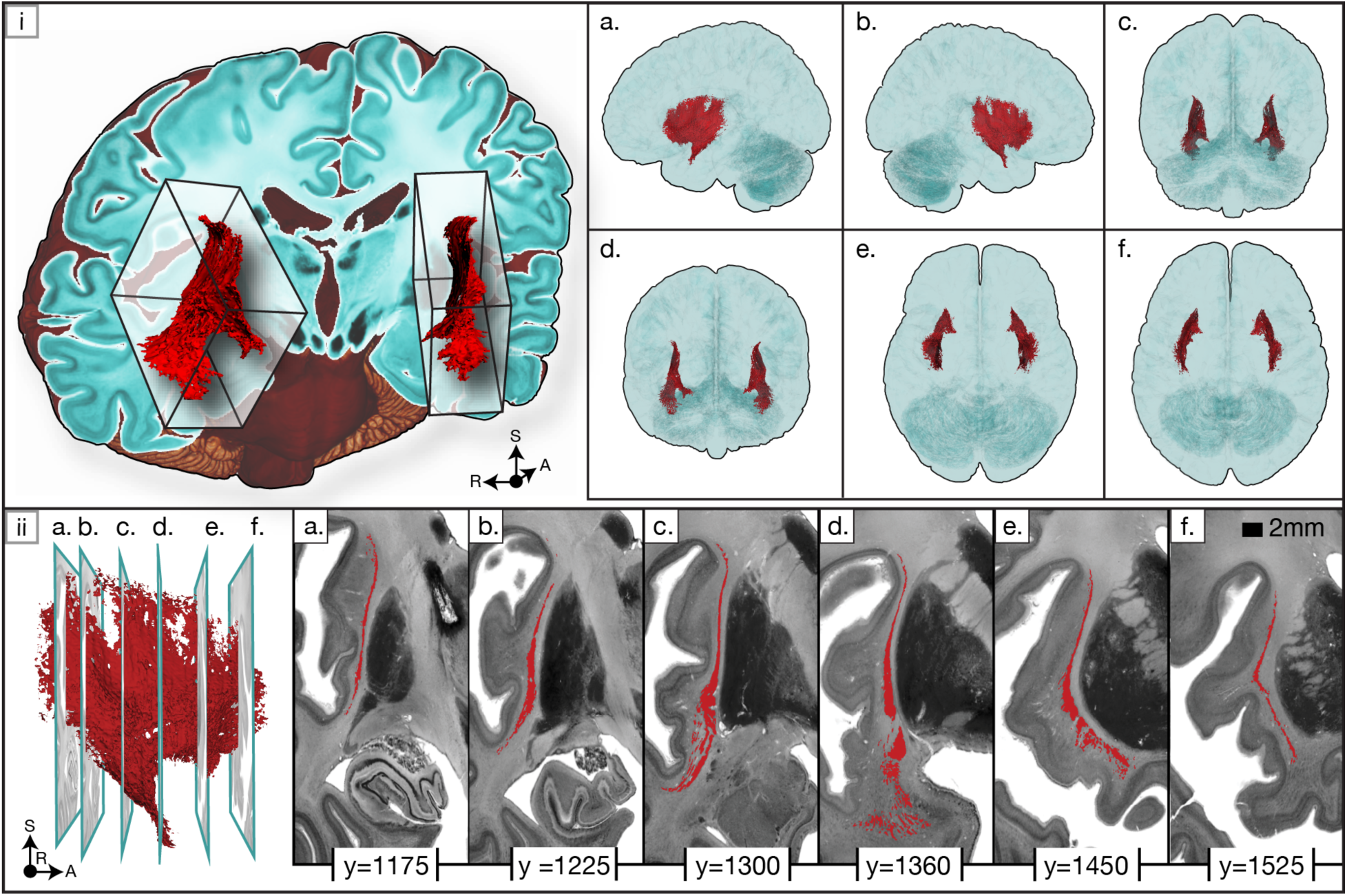
The histological gold standard in three-dimensional and two-dimensional views. *[i*] Left: Both claustra of the gold-standard model (red) are shown within the histological BigBrain dataset. Inset shows oriented bounding boxes (OBB) from anterior view, revealing oblique orientation relative to cardinal axes. Right: Six canonical views (left, right, posterior, anterior, inferior, superior) highlight the claustrum’s shape and position within the brain. [*ii*] Left: Lateral view of the right claustrum with six coronal slice positions indicated (a–f). Right: Segmentations of the corresponding slices are shown in coronal view (BigBrain coordinates provided). The claustrum shows substantial anterior–posterior variability: aligning with insular cortex posteriorly (a–b), fragmenting into ventral “puddles” mid-depth (c–d), and curving around putamen anteriorly (e–f).

The two greatest segmentation challenges at 100μm compared to 20µm BigBrain histology arose from features that are difficult to resolve even at cellular resolution: we could not detect tiny islands abutting the piriform cortex near the terminal zone of the lateral olfactory tract^5,55^, and some boundaries with the amygdaloid complex in the anterior ventral claustrum were ambiguous^7,56^. Because the model did not reveal a clear structural basis for delineating putative claustral subsections, and it is unclear whether such subdivisions can be reliably distinguished on cytoarchitectural grounds alone^57^, we adopted the rhinal sulcus as a practical heuristic to separate dorsal from ventral claustrum^58^.

BigBrain’s continuous reconstruction and our full segmentation enable more precise morphometry than interpolation across sectioned histology. Bilateral centres of mass were symmetric at approximately ±32mm from midline, 1mm posterior to the anterior commissure, and 5-6mm inferior to the anterior commissure line. The principal axes showed an oblique trajectory, with anterior (∼40°) and inferior (∼50°) deviation relative to canonical neuroanatomical planes. Three-dimensional measurements averaged across hemispheres are presented in **Table 1** (hemisphere-specific results in **Extended Data Table 1**). Total claustrum volume was 2536.02mm³ (left: 1325.58mm³; right: 1210.44mm³), approximately 0.13% of the total brain volume, including cerebellum and ventricular CSF. Maximal axis-aligned extents measured 28.35mm mediolaterally, 53.45mm anteroposteriorly, and 55.45mm superoinferiorly. Shape descriptors indicated low roundness and high flatness, consistent with an elongated, planar structure.

**Table 1.** Morphological metrics for the gold standard, the gold standard downsampled to three acquired MRI resolutions (thresholded at 50%), and MRI datasets. Values are averaged across hemispheres. For the gold standard and downsampled gold standards (one brain), bracketed values indicate inter-hemispheric differences and should not be interpreted as a true standard deviation. For MRI datasets (each n=10), bracketed values indicate standard deviation. See **Extended Data Table 1** for hemisphere-specific results.

|  | Gold standard | Downsampled gold standards |  |  | MRI datasets |  |  |
| --- | --- | --- | --- | --- | --- | --- | --- |
| Resolution | 100µm | 0.5mm | 0.7mm | 1.0mm | 0.5mm | 0.7mm | 1.0mm |
| *Three-dimensional* |  |  |  |  |  |  |  |
| **Volume (mm<sup>3</sup>)** | 1268.01<br>(81.42) | 1042.12<br>(101.65) | 1030.54<br>(106.96) | 864.00<br>(77.78) | 1459.14<br>(183.20) | 1371.33<br>(126.61) | 1318.80<br>(400.38) |
| **Maximal x extent (mm)**<br>(mediolateral) | 28.35<br>(2.90) | 22.00<br>(2.12) | 21.00<br>(1.98) | 19.00<br>(1.41) | 18.65<br>(1.53) | 16.91<br>(1.23) | 15.35<br>(1.95) |
| **Maximal y extent (mm)**<br>(anteroposterior) | 53.45<br>(4.03) | 45.00<br>(0.71) | 44.80<br>(0.99) | 40.50<br>(0.71) | 47.83<br>(3.46) | 45.71<br>(3.37) | 39.75<br>(4.40) |
| **Maximal z extent (mm)**<br>(inferosuperior) | 55.45<br>(2.19) | 51.75<br>(0.35) | 49.70<br>(0.99) | 41.00<br>(0.00) | 36.42<br>(2.25) | 35.49<br>(3.74) | 33.00<br>(2.38) |
| **OBB x'** | 24.41<br>(2.30) | 18.58<br>(2.55) | 17.89<br>(2.22) | 16.71<br>(0.15) | 15.74<br>(1.42) | 14.52<br>(1.24) | 11.96<br>(1.37) |
| **OBB y'** | 47.35<br>(0.49) | 45.30<br>(0.11) | 45.05<br>(0.90) | 36.56<br>(3.65) | 36.59<br>(1.94) | 34.85<br>(3.59) | 32.83<br>(2.02) |
| **OBB z'** | 57.26<br>(1.62) | 50.35<br>(0.73) | 50.19<br>(0.93) | 44.50<br>(5.10) | 48.16<br>(3.18) | 46.61<br>(3.01) | 41.10<br>(4.24) |
| **Roundness** | 0.08<br>(0.00) | 0.20<br>(0.00) | 0.23<br>(0.00) | 0.31<br>(0.01) | 0.23<br>(0.01) | 0.27<br>(0.01) | 0.35<br>(0.02) |
| **Flatness** | 3.58<br>(0.62) | 3.46<br>(0.72) | 3.39<br>(0.72) | 3.17<br>(0.75) | 3.51<br>(0.43) | 3.46<br>(0.33) | 3.80<br>(0.45) |
| *Two-dimensional* |  |  |  |  |  |  |  |
| **Mean thickness, total voxel span (mm)** | 0.97<br>(0.60) | 1.15<br>(0.48) | 1.25<br>(0.46) | 1.44<br>(0.34) | 1.69<br>(0.50) | 1.75<br>(0.51) | 2.14<br>(0.49) |
| **Mean thickness, contiguous voxels (mm)** | 0.48<br>(0.17) | 0.97<br>(0.32) | 1.10<br>(0.32) | 1.39<br>(0.31) | 1.61<br>(0.43) | 1.72<br>(0.49) | 2.14<br>(0.49) |

To characterise the claustrum’s thinness and ventral fragmentation, we computed two-dimensional (slice-wise) thickness metrics that mitigate potential overestimation of maximal three-dimensional extents. Across coronal slices, the mean span of mediolateral thickness was 1.21mm±1.39mm, whereas the thickness of contiguous voxels was just 0.56mm±0.52mm. Discrepancies between these measures occurred in >90% of slices and exceeded a twofold difference in 40%, indicating interruption of white matter fibres, primarily in regions containing ventral “puddles” (**Fig. 3**). All coronal slices contained submillimetre spans, and 85% contained at least one location only one voxel thick (100 µm). Thickness maps projected along orthogonal planes revealed a counterintuitive pattern: although the dorsal claustrum forms a narrow sheet, it contains a relatively cohesive central ‘core’, whereas the ventral claustrum, despite its broad mediolateral span, contains fewer claustrum voxels due to punctuation by white matter (**Fig. 4**).

**Fig. 3.**
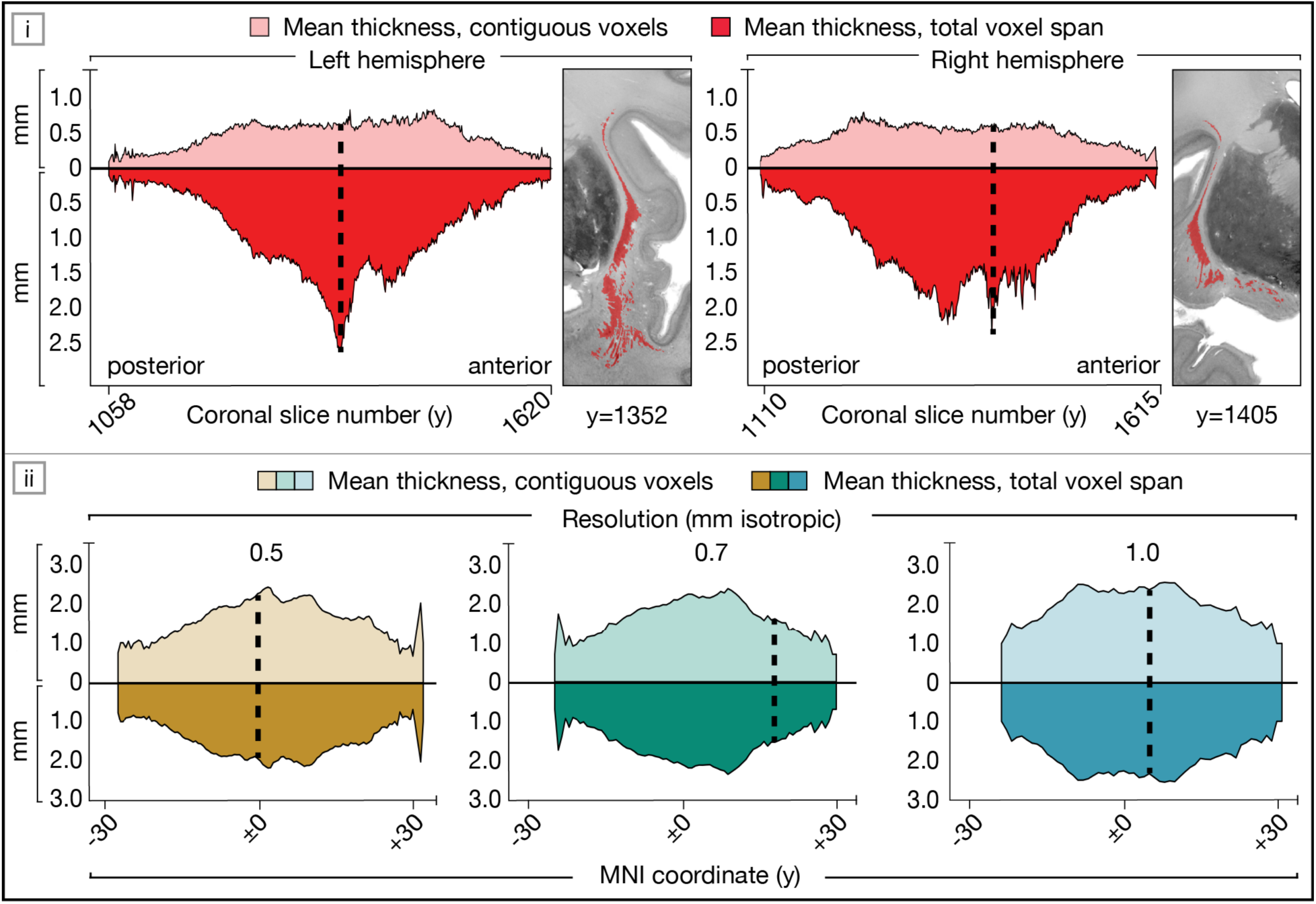
Slice-wise thickness metrics in the histological gold standard and MRI. In each silhouette plot, mean thickness of contiguous voxels (upper, pastel) and total voxel span (lower, solid) are mirrored around the horizontal axis. The dashed black line indicates the coronal slice with maximal discrepancy between measures. [i] Gold standard thickness profiles shown separately by hemisphere, with insets highlighting slices of maximal discrepancy. [ii] Mean thickness profiles from three MRI datasets, averaged across participants and hemispheres. Discrepant ratios in the gold standard reflect ventral “puddles” perforated by white matter; these are absent in MRI where ratios approach unity.

**Fig. 4.**
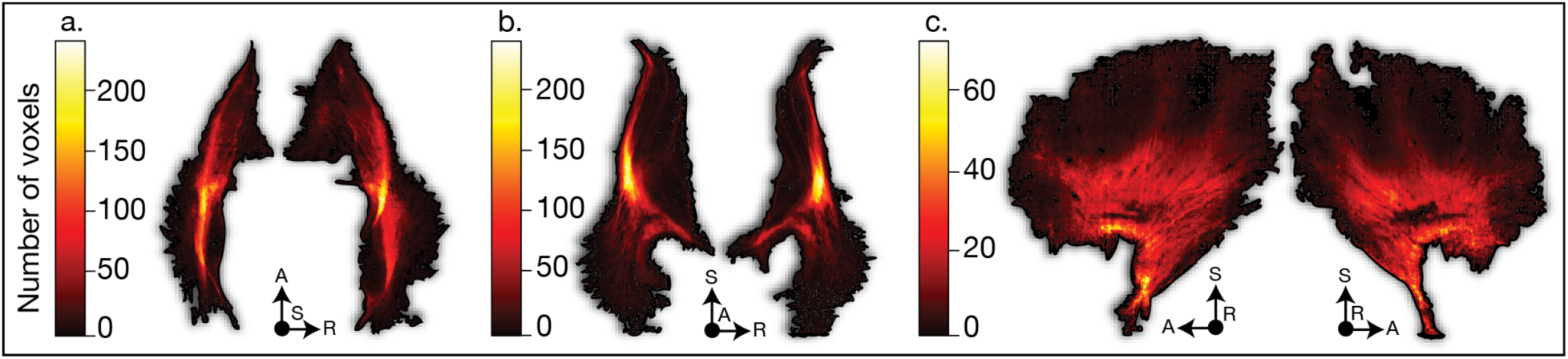
Two-dimensional thickness map of the histological gold standard. To visualise claustral thickness, the three-dimensional gold standard was projected into two dimensions, with colour indicating voxel count (dark = low count, light = high count). Projections are shown in the (a) axial, (b) coronal, and (c) sagittal planes for the left and right claustra, respectively. A truncated scale is employed for the sagittal view, required for visual distinction. The thickest regions, represented by the bright yellow and red “core,” are comparable to the claustrum’s appearance in submillimetre MRI (see Fig. 8) and reliably identified across participants across MRI resolutions (see **Extended Data Fig. 1**).

### Downsampling systematically alters claustral geometry

To assess how spatial resolution affects claustrum morphometry, we downsampled the histological gold standard to MRI-like isotropic resolutions (from 0.4–2.0mm, in 0.1mm increments), and across binarisation thresholds (0.2–0.8) reflecting liberal vs. conservative segmentation style. Resolution exerted heterogeneous effects on claustral geometry, but the resulting degradation was predictable, with resolution alone explaining more than 93% of variance across all eight metrics (**Fig. 5**). Volume was largely insensitive to voxel size (p=0.18). However, all three maximal axis-aligned extents decreased systematically with coarser resolution (all p_FDR_< 0.01), shrinking by 0.16mm mediolaterally, 0.19mm anteroposteriorly, and 0.40mm superoinferiorly per 0.1mm increase in resolution. Roundness increased at lower resolution, while flatness remained stable except at the lowest resolutions and highest thresholds, where it dropped sharply, reflecting a shift toward a less elongated and planar claustral geometry (both p_FDR_<0.01). Consistent with partial voluming, both total voxel span and contiguous voxel thickness increased at coarser resolution (p_FDR_<0.01).

**Fig. 5.**
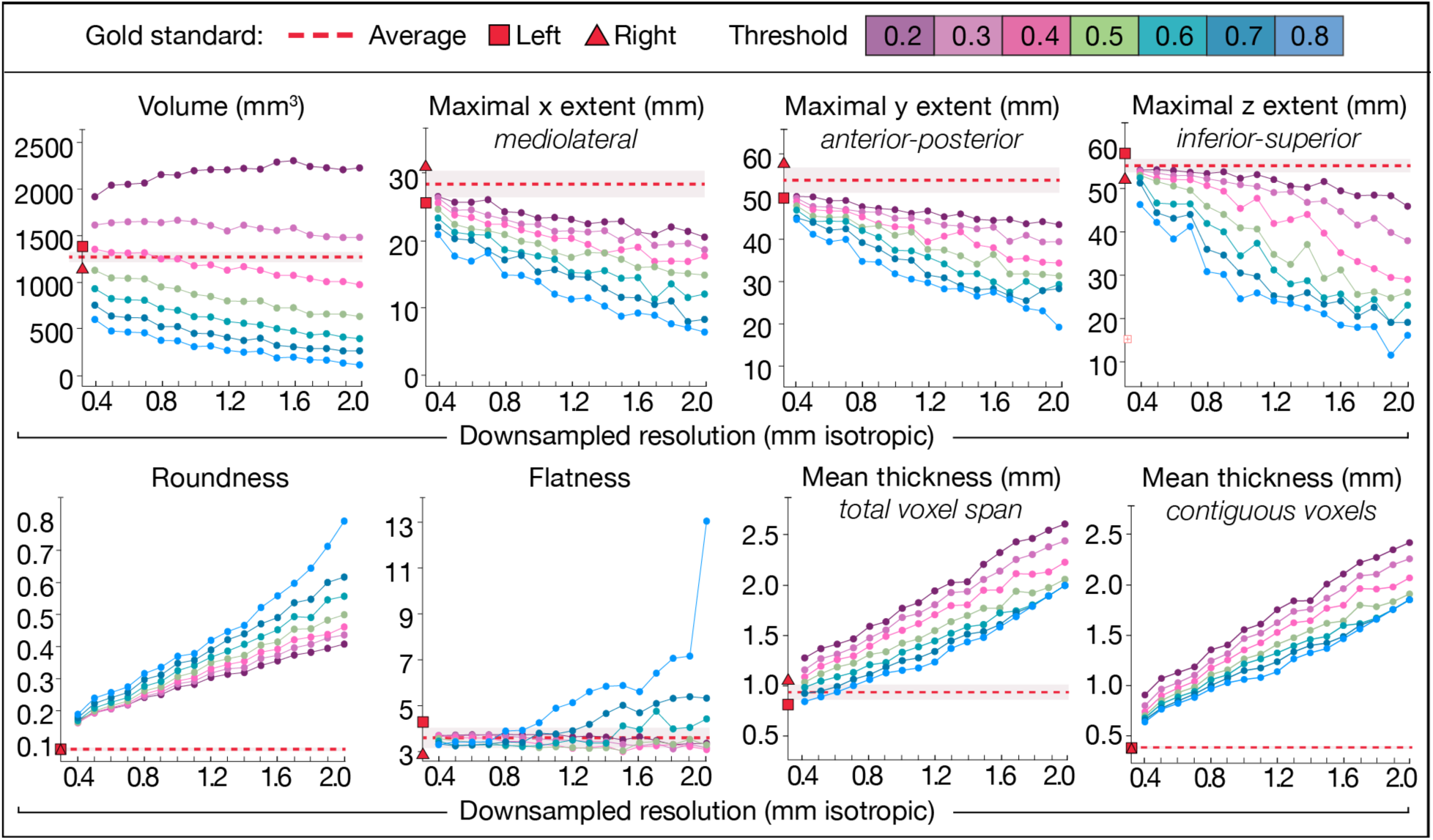
Downsampling of the histological gold standard to various MRI-like resolutions, at different thresholds. Downsampled estimates of the eight morphometric measurements, averaged across hemispheres, to resolutions of 0.4-2.0mm. The gold standard’s measurements are shown in red (dashed line = average, square = left hemisphere, triangle = right hemisphere). Each coloured line represents a different binarisation threshold (0.2-0.8).

### MRI partially captures claustral anatomy

In all three ultra-high field T1-weighted MRI datasets (0.5, 0.7, and 1.0mm isotropic resolutions), the claustrum appeared hypointense with adequate contrast-to-noise ratio (CNR) relative to surrounding white matter (0.5mm = 4.42±0.52, 0.7mm = 3.29±0.39, 1.0mm = 2.78±0.60). Though raters reported lower confidence in boundary delineation compared to histology, inter-rater agreement remained high (all DSC>0.9, see **Supplementary Fig. 1**). The claustrum’s proportion of intracranial volume was consistent across datasets: 0.27%± 0.04 at 0.5mm, 0.26%±0.04 at 0.7mm, and 0.25%±0.09 at 1.0mm, and centres of mass were likewise stable (maximum difference 1.67mm; **Supplementary Table 1**). However, only the 0.5mm dataset enabled unambiguous differentiation in all participants, with manually-drawn claustrum segmentations sometimes abutting but never overlapping adjacent cortical or subcortical structures, despite partial voluming with white matter (**Fig. 6**).

**Fig. 6.**
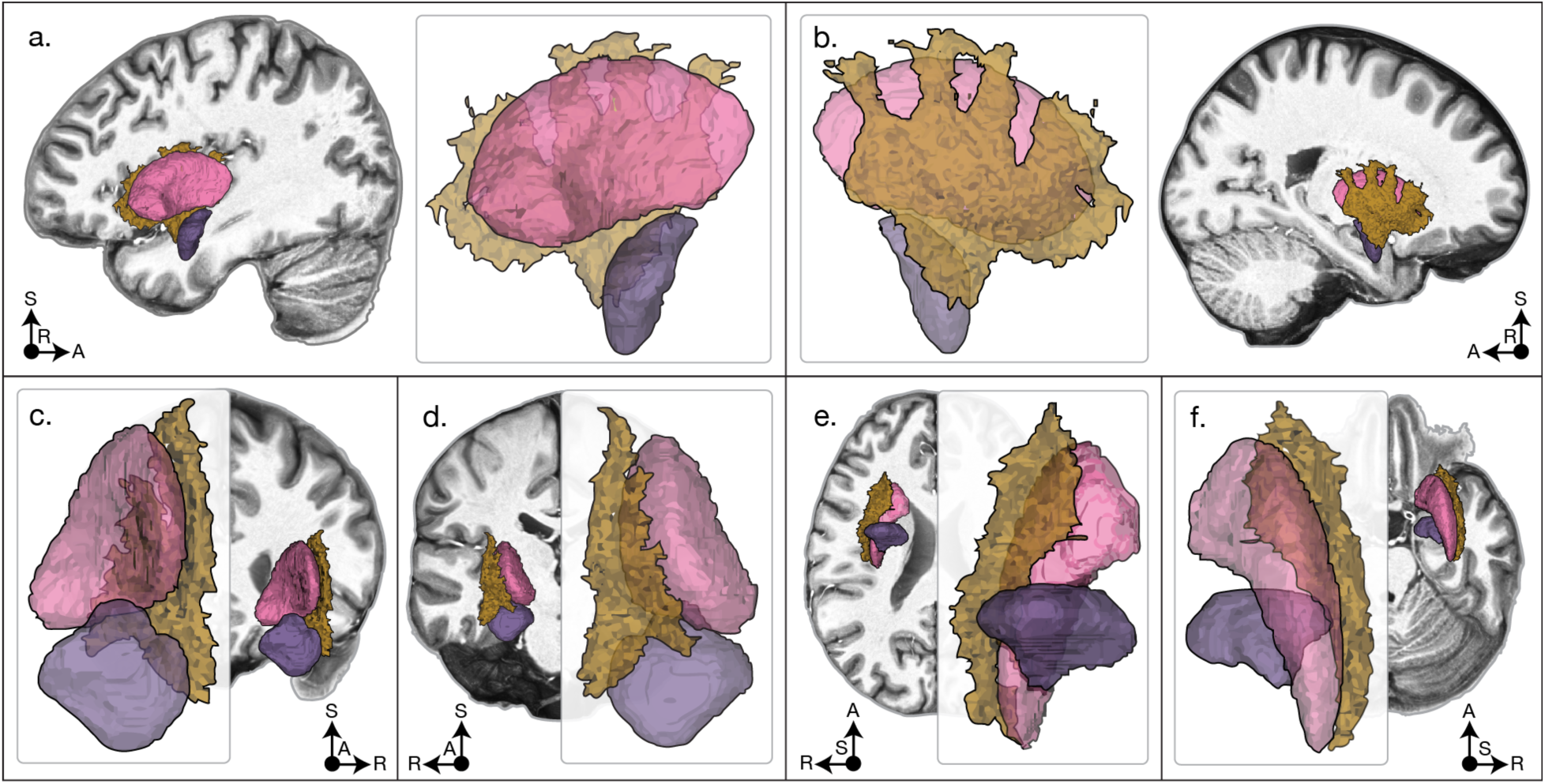
MRI capture of the claustrum at 0.5mm isotropic resolution. The manually segmented claustrum (yellow) from a representative subject in the 0.5mm dataset is shown in six canonical views (a–f: left, right, posterior, anterior, inferior, and superior), alongside the putamen (pink) and amygdala (purple) as labelled by the Xiao atlas (2019). The background slice in each panel is the nearest slice that does not contain claustral voxels. Among the MRI datasets, only the 0.5mm resolution reliably and unambiguously distinguishes all three structures.

Comparison of claustral morphometry across MRI datasets is visualised in **Fig. 7** and quantified in **Extended Data Table 2**. As in the downsampling simulation, claustrum volume was not significantly affected by resolution (p_FDR_=0.25), but all other metrics showed significant resolution effects, from which two patterns emerged. First, 1.0mm showed systematically greater divergence from the submillimetric datasets: *post hoc* tests showed that the 0.5mm and 0.7mm datasets differed only occasionally (3 of 7 significant comparisons), whereas the 1.0mm dataset differed from both submillimetric datasets across all significant metrics. Second, measurements were generally less stable at 1.0mm, with volume exhibiting markedly high instability (CV=0.31; see **Supplementary Table 2**). To support comparable claustrum findings across studies, we propose practical reporting standards (**Supplementary Note 1**).

**Fig. 7.**
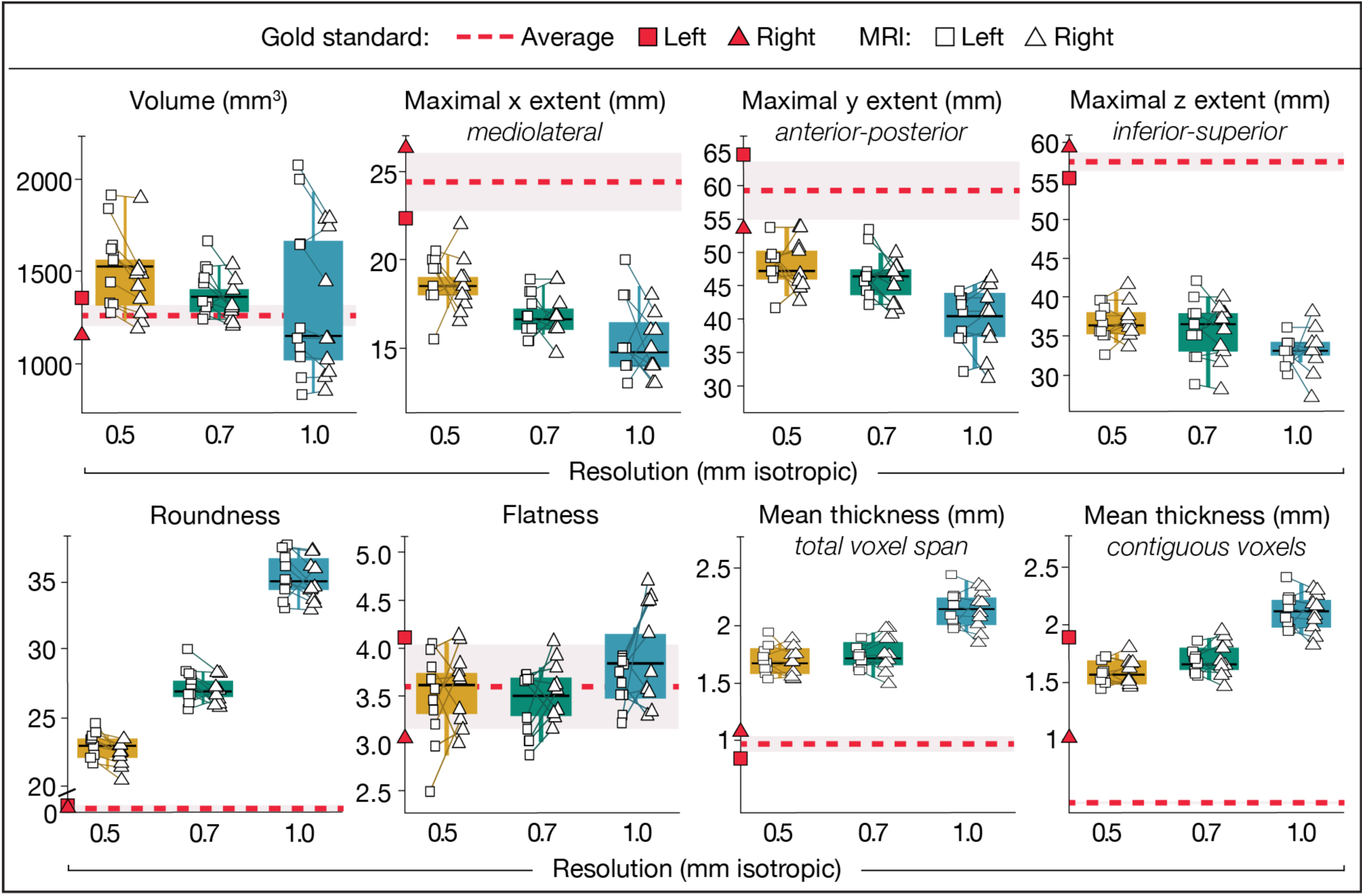
Resolution-dependent effects on MRI-derived claustrum morphometry. Estimates of the eight morphometric measurements are shown for MRI datasets. Boxplots depict values averaged across hemispheres; solid horizontal lines depict the median. Squares and triangles represent the left and right hemisphere values, respectively. For comparison, the gold standard measurements are indicated in red (dashed line = average). All morphometric measures showed clear resolution-dependent changes except volume, reflecting the volume paradox: mediolateral thickening combined with anteroposterior and dorsoventral truncation produced similar total volumes despite fundamentally altered morphology.

### MRI capture diverges from the histological gold standard

As anticipated, direct comparison between MRI segmentations and the histological gold standard revealed substantial deviations across most morphometric measures (**Fig. 7 and Extended Data Table 3**). Both submillimetre MRI datasets overestimated claustum volume, most prominently at 0.5mm; the 1.0mm dataset did not differ significantly. Flatness was the sole metric that remained stable across all resolutions, reflecting proportional shrinkage along the anteroposterior and superoinferior axes. All other measures showed significant resolution-dependent deviations that increased with coarser resolution. Two-dimensional thickness estimates showed especially large discrepancies (total span +74–121%; contiguous thickness +234–344%) (**Fig. 3**). Fidelity was poorest in the middle third of the anteroposterior axis where the ventral claustrum broadens and fragments into “puddles”: the gold standard’s span-to-contiguous-thickness ratio (2.76) collapsed to near-unity at all MRI resolutions, indicating near-complete loss of anatomical detail (**Fig 3. and Supplementary Table 3**).

Spatial agreement between MRI and the gold standard indicated poor correspondence (**Extended Data Table 4**). Dice coefficients were uniformly low (DSC 0.37–0.40), while Hausdorff Distances were high and increased with coarser resolution (HD 9.49mm–13.05mm). When boundary uncertainty was accommodated using adjusted metrics (dilated DSC and balanced average HD), spatial agreement improved substantially, indicating that MRI-histology discrepancies reflected boundary imprecision rather than gross mislocalisation (**Fig. 8i**).

**Fig. 8.**
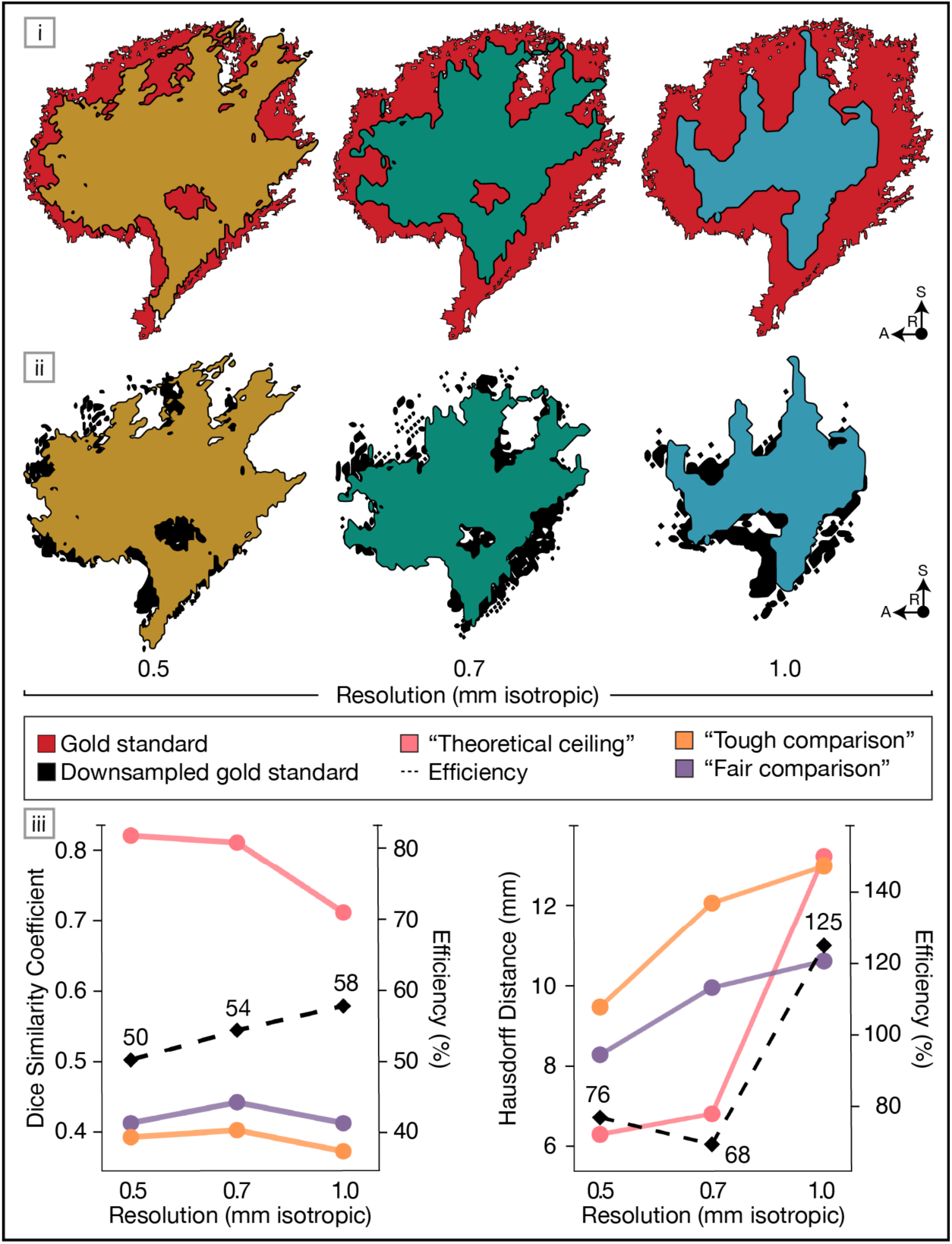
Spatial agreement between histological gold standard(s) and MRI. [*i–ii*] Overlap of MRI segmentations (colour) with either the histological gold standard (red; “tough comparison”) or the resolution-matched downsampled gold standard binarised at a 50% threshold (black; “fair comparison”). Lateral views of the left hemisphere are shown for a representative participant with median claustrum volume. Across resolutions, the central core of the claustrum is consistently recovered, whereas peripheral boundaries are progressively lost. [*iii*] Agreement is quantified using Dice similarity coefficient (left) and Hausdorff Distance (right). Orange lines represent MRI performance against the histological gold standard (tough comparison), and purple lines represent MRI performance against the resolution-matched downsampled gold standard (fair comparison). Pink lines show the theoretical ceiling (i.e., downsampled vs. full-resolution gold standard), and dashed black lines indicate MRI efficiency (the proportion of achievable performance attained at each resolution). MRI captures roughly half of the maximal possible volumetric overlap (DSC) and 68–76% of attainable boundary precision (HD) at submillimetre resolutions. At 1.0mm, reduced ceiling performance (pink) produces inflated HD efficiency despite poorer absolute boundary accuracy. Exact values provide **Supplementary Table 4.**

### MRI approaches the sampling ceiling of resolution-matched downsampling

Finally, each MRI dataset was evaluated against its resolution-matched downsampled gold standard binarised at a 50% threshold, which we took as the theoretical maximum detail recoverable at a given resolution (**Extended Data Table 5**). Again, MRI consistently overestimated volume, albeit most strongly at 1.0 mm (+53.34%). Maximal mediolateral and superoinferior extents were truncated across all MRI datasets, while anteroposterior extent was significantly longer at 0.5 mm (+6.29%) but comparable at 0.7 mm and 1.0 mm. Roundness showed mild inflation, and flatness remained stable except for an increase at 1.0 mm (+22.42%). Slice-wise thickness was consistently overestimated, with the largest deviations observed at coarser resolutions (total span +40–48%; contiguous thickness +53–67%).

In addition to spatial agreement metrics, we computed ‘efficiency’ as the proportion of agreement attainable given the downsampled gold standard’s inherent ceiling (**Supplementary Table 4)**. DSC was low and resolution-invariant (0.41–0.44), corresponding to 50.00% to 57.75% of theoretically achievable volumetric overlap. HD ranged from 8.29 mm to 10.65 mm, representing 68.14% to 75.75% of attainable boundary precision at submillimetre resolutions; at 1.0 mm, however, the downsampled gold standard exhibited such poor boundary definition that MRI performance nominally exceeded the ceiling (124.69%), underscoring that claustral boundaries are poorly represented at ‘conventional’ resolution (**Fig. 8ii**).

### Inter-individual variability, hemispheric asymmetry, and sex differences

Despite morphometric distortion, MRI is arguably the best tool for studying claustral variation *in vivo*. Thus, we pooled the three MRI datasets to explore individual variability, hemispheric asymmetry and sex differences, while acknowledging inherent measurement limitations.

#### Individual variability

A probabilistic overlay constructed from all 30 MRI segmentations revealed high spatial agreement in the dorsal “core” of the claustrum and progressively lower agreement toward the ventral extent (**Extended Data Fig. 1**).

#### Hemisphere differences

In the pooled MRI sample, the right claustrum was significantly larger in volume (*p*_FDR_<0.01, *d*=0.91) and exhibited greater flatness (*p*_FDR_=0.03, *d*=0.47), whereas the left claustrum was more round (*p*_FDR_<0.01, *d*=0.75). In the 0.5mm dataset, participant-level asymmetry indices confirmed significant hemispheric asymmetry in volume (AI=-0.036, *p*_FDR_=0.019) and roundness (AI=0.021, pFDR=0.014). Asymmetry patterns were consistent across resolutions.

#### Sex differences

Comparison between sexes revealed no significant differences on any morphometric measure, between or across hemispheres; likewise, controlling for intracranial volume (ICV) revealed no significant effects of sex or ICV.

## DISCUSSION

Despite decades of interest in claustral function, direct study in living humans has remained elusive because of pessimism that MRI’s spatial resolution is inadequate to capture the structure’s unusual geometry. Here, we ask whether MRI can resolve the human claustrum with sufficient fidelity to support *in vivo* investigation, by characterising its anatomy using complementary approaches: high-resolution histology (a 100µm BigBrain-derived gold standard model) and ultra-high field MRI (7-Tesla datasets at 0.5mm, 0.7mm, and 1.0mm isotropic resolution). Through systematic comparison of these modalities and resolution-matched simulations, we establish what each captures of claustral structure, quantify the limits of *in vivo* imaging, and provide a histology-based benchmark that lays the foundation for next-generation studies of human connectivity and function.

The BigBrain-derived histological gold standard model provides the first continuous three-dimensional reconstruction of the human claustrum derived directly from serial histological sections, without statistical interpolation. This publicly-available, interactive model enables appreciation of claustrum size, complexity, and anatomical relationships in a way that traditional illustrations and photographs cannot (**Fig. 2**). The model also highlights striking architectural contrasts: although the claustrum is often only a few hundred microns thick, it spans more than 5cm anteroposteriorly and superoinferiorly, with a total bilateral volume twice that of the substantia nigra and approaching three-quarters that of the amygdala^59^. Its large extent belies common descriptions of the claustrum as a “tiny” nucleus^60^, but its thinness and undulation helps explain why many have assumed it to be beyond the reach of conventional MRI.

The gold standard resoundingly accords with qualitative descriptions and illustrations of classical anatomical literature^5–7^. Direct comparison is limited by sparse reporting of quantitative metrics and pronounced methodological differences, including fixed versus fresh tissue and varying conversion factors. Our bilateral volume lies at the upper end of published histological estimates (**Fig. 1**): only one reports a slightly higher value^61^, whereas five report smaller volumes^33,48,55,62,63^. Only one prior study quantified extents and reported a substantially shorter anteroposterior and dorsoventral span but larger mediolateral span, likely reflecting coarser sampling^48^. We attribute the comparatively larger measurements in our gold standard to complete manual segmentation of the entire claustrum, made possible by BigBrain’s high tissue integrity and visual contrast.

Downsampling the gold standard isolates resolution-driven distortion and establishes a critical interpretive guardrail for MRI: assuming cell-stained histology affords equal or better identification of claustral tissue than voxelised MRI, any anatomical feature that disappears in downsampling simulations may not be reliably detected in MRI at the corresponding resolution, regardless of apparent visualisation. As expected, discrepancies were greatest at the lowest resolutions and most conservative thresholds (**Fig. 5**), but even the highest MRI-like resolution and most liberal thresholds fundamentally altered claustral morphology and struggled to preserve the ventral claustrum. Simultaneously, the superoinferior extent was disproportionately truncated—reflecting loss of the ventral portion extending into the temporal lobe—yet in regions that remained resolved, contiguous slice-wise thickness inflated as small gaps were bridged and isolated “puddles’’ merged. Roundness increased as thin edges disappeared, while flatness remained stable until the coarsest resolutions artefactually inflated mediolateral thickness and eliminated sheet-like geometry. This progressive degradation explains why even high-resolution MRI studies typically visualise the claustrum as a simplified ribbon lacking ventral extension.

A notable consequence of this degradation was a “volume paradox”: despite marked truncation of the claustrum’s anteroposterior and dorsoventral extents, total volume remained statistically stable across resolutions because the reduction in extents was offset by partial-volume inflation of mediolateral thickness. Thus, volume stability does not indicate preserved anatomy but rather that volume is an insufficient descriptor of claustral morphology. This phenomenon is also described in other thin structures where boundary voxels disproportionately influence total volume^64,65^. Further, downsampling to 1.0mm with a 50% threshold—optimistically representing the most common *in vivo* MRI resolution and a typical ‘majority-vote’ segmentation approach—produced substantial divergence from histological ‘reality’ across nearly all morphometric measures. This implies that MRI at conventional resolution characterises substantial resolution artefacts alongside anatomy, raising questions about interpretation of the extant literature.

We next quantified the extent to which the claustrum, as illuminated by the gold standard model, can be captured *in vivo.* Three ultra-high field datasets established that the claustrum can be (at least partially) identified using standard whole-brain MP2RAGE protocols feasible at most 7-Tesla centres, requiring no specialized contrast^66^. Partial volume effects, evident as intermediate signal intensities at tissue interfaces and loss of anatomical detail, were apparent at all resolutions but more pronounced as voxel size increased; this is expected given that increasing isotropic spatial resolution from 1.0mm to 0.7mm and 0.5mm decreases volume by factors of approximately 3 and 8 (∼1000nL to 343nL and 125nL)^67^. The 0.5mm dataset uniquely separated the claustrum from surrounding structures (**Fig 6**), though at all resolutions, at least one participant exhibited some degree of apparent ventral “dropout”, almost certainly artefactual rather than true absence given histology’s consistent demonstration of ventral claustrum^68^, albeit with some shape and density variability^6^. No aspect of the claustrum exhibited markedly different contrast properties despite known variation in neuronal density^33^.

As in downsampling, MRI showed paradoxical volume stability across datasets despite shape changes (**Fig. 7**). This stability contrasts sharply with dramatic between-study variability in the literature, with higher-resolution studies tending to report larger volumes (**Fig. 1**). The most likely explanation is segmentation style: the claustrum’s extreme thinness makes measurements highly sensitive to boundary decisions. Illustratively, though we implemented the segmentation protocol of Kang and colleagues^69^ on resolution-matched data, we obtained bilateral volumes ∼20% smaller than theirs. Likewise, independent groups^51,52^ segmenting the same *ex vivo* brain^70^ differed by 18% in one hemisphere. Rigorous standardization of manual segmentation or automated algorithms are needed; we recommend reporting standards to make claustrum findings interpretable and comparable across studies (**Supplementary Note 1**).

While decades of MRI-based claustrum research have acknowledged potential limitations, none have tested these assumptions against a histological reference, creating an evidence base of uncertain reliability. Direct comparison to the gold standard revealed poor spatial agreement (**Fig. 8i**) and substantial deviations across all morphometric measures (**Fig. 3**, **Fig. 7**), except paradoxically-stable volume. However, all resolutions showed a “parochial” detection pattern: MRI reliably captured thick core regions while losing thin peripheral features. The preserved core corresponded to thick mid-dorsal regions in the histological map (**Fig. 4**), whereas thin boundaries escaped detection, including much of the ventral claustrum but also superior dorsal aspects where the claustrum bends over the putamen. When boundary uncertainty was accommodated using adjusted metrics appropriate for thin structures (dDSC and baHD), overlap was reasonable at both submillimetre resolutions.

Importantly, the claustrum’s thickest dorsal portions account for most claustral density and volume^33^, and house primary connectivity to sensorimotor and frontal association cortices^2,71,72^, grounding distinct hypotheses of claustral function^3,73,74^. Other subcortical research has succeeded under such constraints: hippocampal studies focus on CA1 and dentate gyrus while accepting poor CA2/CA3 resolution^75^, and substantia nigra work routinely targets ventral tiers despite dorsal detection failures^76^. Such constraints have not stymied progress but have prompted greater anatomical precision, more targeted hypotheses, and appropriately cautious interpretation: a mature scientific approach the claustrum field now requires.

Next, we asked if MRI’s limitations reflect resolution constraints or other technical factors, so compared each MRI dataset to its resolution-matched downsampled gold standard binarized at 50% threshold (**Fig 8ii**). This “fair comparison” isolates spatial sampling effects from other potential sources of discrepancy such as the sensitivity of MRI contrast to histologically-determined cell density. MRI distorts claustral anatomy through mechanisms largely, but not entirely, explained by spatial sampling. Submillimetre MRI achieved approximately half of theoretically achievable volumetric overlap and the majority of attainable boundary precision (**Fig 8iii**). Importantly, our MRI datasets were not optimised for claustral nor subcortical capture, suggesting the 25-50% efficiency shortfall may reflect correctable technical factors rather than fundamental limits. Potential optimization could include tuning echo time for claustral contrast, testing slice-plane angulation relative to the insular sheet, exploring modest anisotropy as used for other small subcortical structures^77^, and testing alternative contrasts that may enhance capsule boundaries and have been useful to automated segmentation efforts^52,78^.

Cross-modal contrast differences likely also contribute. Two independent annotations^51,52^ of the same 100µm *ex vivo* 7-Tesla MRI dataset^70^—one using multi-planar, multi-rater segmentation with union smoothing, the other a sparse single-rater coronal approach with interpolation—yielded deviations from the gold standard, including inflated volumes, truncated extents, higher roundness, and span-to-contiguous ratios near unity (**Supplementary Table 5)**. These discrepancies mirror those observed in our *in vivo* MRI and downsampling analyses, suggesting they may arise from MRI contrast properties rather than spatial sampling alone. Thus, though technical optimization may improve claustral imaging within existing 7-Tesla infrastructure, cell-stained histology remains the necessary reference for precise anatomical characterisation.

How do these results bear on studies of claustral function? Even at high- and ultra-high field, typical fMRI resolution (∼1.5-3.0mm) falls far below the submillimetre resolution required for reliable claustral localisation (i.e., voxels containing predominantly claustral tissue). Echo-planar imaging further degrades effective resolution through T2* blurring, susceptibility-induced distortion, and corrective resampling artefacts^79^, with additional confounds likely arising from insular perforators of the middle cerebral artery and venous drainage^80^. Yet anatomical invisibility does not preclude functional detection: voxels containing claustral tissue can generate measurable BOLD signal despite partial volume dilution^81^. Two 7-Tesla fMRI studies provide proof-of-concept: Coates and colleagues detected task-evoked responses at 1.34×1.34×0.8mm resolution^17^, and Krimmel and colleagues recovered claustral resting-state correlations at 1.5mm isotropic resolution, explicitly addressing contamination from the insula and putamen^82^.

The histological gold standard can guide functional investigation by providing an anatomical prior, ensuring invisibility in structural MRI does not preclude detection via fMRI. Warping the gold standard atlas into subject space allows, for example, principled seed placement in resting-state fMRI and interpretation of apparent white matter activations in task fMRI. To better account for interindividual variability and bridge histology and MRI, we also provide a cross-modality probabilistic atlas integrating the gold standard, the 0.5mm 7-Tesla dataset (n=10), and two high-resolution MRI segmentations made publicly available by Mauri and colleagues^52^, one *ex vivo* at 100µm^70^ and one *in vivo* at 250µm^83^. See **Supplementary Note 2** for probabilistic atlas description.

Finally, exploratory analyses across the three MRI datasets revealed reproducible patterns despite demonstrated limitations of *in vivo* resolution. The probabilistic overlay showed highest spatial agreement in the dorsal midsection, with increasing variability toward the superior, anterior, and ventral periphery where partial voluming was most pronounced (**Extended Data Fig. 10**), underscoring that atlases for MRI must be derived at substantially higher resolution^84^ (see **Supplementary Note 2**). Hemispheric asymmetry was modest but consistent: the right claustrum appeared larger and flatter, the left smaller and rounder. This contrasts with the leftward bias in the histological gold standard. Prior MRI reports are mixed: some report rightward trends in adults^12,69,85,86^, significant rightward effects in adolescent males^87^ and neonates^88^, and others report nonsignificant^89^ or significant leftward effects^11,90^. Two independent segmentations of the same 100µm *ex vivo* MRI^70^ also found a larger right claustrum^51,52^. No significant sex differences emerged, consistent with most prior studies showing absent effects^85^ or higher male volumes that disappear after ICV adjustment^69^ or do not reach significance^12^, though one study found higher female volumes after ICV adjustment^86^, and subtle tissue-composition differences have been reported^89,91^. Collectively, these results support pooling sexes and modelling hemispheres separately to maximise statistical power.

Our approach has several limitations alongside strengths advancing claustrum investigation. The BigBrain-derived gold standard is from a single 65-year-old male brain, limiting assessment of population variability. The 100µm BigBrain smooths some claustral features visible in the 20µm and 1µm versions^68^, but was used due to feasibility, its availability in MNI space, and its widespread adoption. Manual segmentation, including the use of different raters by hemisphere, introduces some subjectivity despite high inter-rater reliability. Nonetheless, pending higher-resolution, multi-donor and multimodal validation, this remains the most complete three-dimensional histological model of the human claustrum available.

Our MRI analysis used three convenience datasets (each n=10) acquired on different Siemens systems with slightly varying protocols, introducing potential site and sequence heterogeneity, though we observed no substantial SNR limitations or distortion artefacts. Participant-related biases cannot be excluded, but demographics were comparable across datasets, and prior subcortical atlasing suggests that morphological estimates stabilise with modest samples (>5)^92–94^. Participants were younger than the BigBrain donor, although current evidence suggests some age effects on claustral morphometry in late adulthood^47,86^. The strengths of this analysis lie in its use of whole-brain sequences that most 7-Tesla centres can implement, and are increasingly available in public datasets.

For more than two decades, the claustrum has been treated as effectively invisible to MRI, pushing human research to the margins of an animal-dominated literature. The present study provides the first continuous three-dimensional histology-based model of the claustrum and the first systematic test of MRI’s ability to capture it. Our results challenge the view, persisting since Crick and Koch, that the claustrum is a tiny nucleus beyond resolve. Submillimetre 7-Tesla MRI recovers more than half of theoretically attainable anatomical detail, reliably capturing the thick dorsal core that comprises most claustral volume and houses major corticoclaustral connectivity hubs^2,33^. Ventral “puddles” remain challenging, yet at 0.5mm isotropic resolution their overall extent is partially preserved, with uncertainty arising from boundary imprecision rather than complete anatomical loss. The current state-of-the-art of *in vivo* MRI permits productive investigation of claustral structure and cautious exploration of its function^13,17^, positioning the field for a new phase of investigation that may answer long-standing questions about the claustrum’s contribution to human cognition.

## METHODS

### Histology

#### Dataset

The BigBrain dataset is an ultra-high resolution digital reconstruction of histological sections from a 65-year-old male with no known neurological or psychiatric conditions at the time of death^53^. It uses a modified silver impregnation method based on Merker’s technique to selectively stain neuronal cell bodies, providing excellent contrast for cytoarchitectural analysis. We selected BigBrain after reviewing publicly available high-resolution digital *ex vivo* datasets, including the MGH atlas^70^ and the Allen Brain Atlas^50^, as the claustrum was the most visually distinct. Given our objective to compare to *in vivo* MRI, we used the 100µm isotropic resolution voxelised version provided in ‘BigBrain3D Volume Data Release 2015’ (https://ftp.bigbrainproject.org/bigbrain-ftp/), aggregating the original 20μm reconstruction of 7,404 histological sections, which includes corrections for tissue shrinkage and is aligned to MNI-ICBM152 2009b symmetric space.

#### Claustrum localisation

Current understanding of human claustrum anatomy is informed by anatomical studies^5–7,95^ and whole-brain histological atlases^49,50^. However, given considerable discordance in boundary illustrations across sources, our delineation prioritised apparent voxel intensity in BigBrain, reflecting the presence of neurons (cell bodies) amongst brighter surroundings (white matter). The delineation is most clearly described in the coronal plane, as inclusive of low intensity (dark) voxels between the insula and putamen, extending into the temporal lobe but excluding the amygdala and piriform cortex.

Within this spatial region, voxels were included as claustral regardless of continuity or cluster size, based on the assumption that all grey matter voxels within these bounds belong to the claustrum. In principle, this approach permits isolated voxels to be labelled as claustral; however, in practice, nearly all included voxels were connected in at least one anatomical plane. We validated this approach by cross-referencing the BigBrain dataset at 20µm and 1µm in-plane resolution^68^, which comprises true cell-stained histology, and confirmed small islands of apparently claustral cells unconnected to the main body in both the dorsal and ventral claustrum.

We segmented the claustrum as a single unified structure. The number, location, and nomenclature of putative subsections have been debated for more than a century^30^ and even modern atlases using similar methods depict markedly different subdivisions^49,50^. In practice, most researchers treat the claustrum as one, using “dorsal” and “ventral” to refer to positions along the superior-inferior axis where morphology markedly differs. We follow this usage and define the ventral region as the portion inferior to the fundus of the rhinal fissure in the middle third of the anteroposterior axis, where the claustrum expands into the temporal lobe and becomes fragmented toward the piriform cortex and amygdala^96^.

#### Segmentation approach

Our approach is illustrated in **Supplementary Fig. 2.** To enable real-time navigation of the massive BigBrain dataset (dimensions: x=1970, y=2330, z=1890), we extracted smaller volumes for each hemisphere encompassing the claustrum and its surrounding structures (dimensions: x_left_ =400-900; x_right_=1000-1500, y=1000-1775, z=400-1000). References to the left and right claustra follow neurological convention.

Manual segmentation was performed in ITK-SNAP^97^ using a Wacom tablet and a one-voxel-sized brush. The right hemisphere was segmented by one rater (SP) and the left by another (NC), following an eight-step quality control process detailed in **Supplementary Note 3**, yielding high inter-rater agreement, with Dice similarity coefficients (DSC) ranging from 0.87 to 0.93 (**Extended Data Fig. 2)**.

Before proceeding with full manual segmentation, we applied various interpolation-based segmentation methods, namely morphological contour^98^, random forest^99^, and SmartInterpol^100^, using default parameters. These methods were tested on a set of 28 consecutive coronal slices in the left hemisphere (y=1335–1365), where the ventral claustrum extends into the temporal lobe. Only every third slice was manually labelled (including the first and last) to serve as input for interpolation. All methods showed more than 15% disagreement (DSC<0.85) relative to manual “ground truth”, particularly struggling with the morphology of the ventral claustrum (**Supplementary Fig. 3**). Additionally, disagreement between the three methods was greater than to manual ground truth (DSC=0.77-0.83). In contrast, two human raters achieve excellent agreement (DSC=0.97) on one representative slice, motivating our decision to proceed with full manual segmentation.

The complete manual segmentation process, including quality control, required approximately 500 hours of labour per hemisphere. Owing to the BigBrain dataset’s unprecedented resolution and the fact that segmentation was conducted manually on each individual two-dimensional slice (with no statistical interpolation), we refer to the resulting three-dimensional reconstruction as the “gold standard” model. To the best of our knowledge, this is the highest-resolution histologically-derived, continuous three-dimensional claustrum model made publicly available.

#### Registration

Due to known subcortical alignment concerns in the original BigBrainSym dataset^42,59^, we ‘re-registered’ BigBrain to an improved MNI-aligned BigBrain^59^ using ANTs SyN, then applied this transformation to the gold standard segmentation using GenericLabel interpolation, for all spatial agreement comparisons to MRI. On the re-registered BigBrain, we also recomputed and compared all morphological metrics, but found only minute differences that did not influence the reported pattern of results; thus, to facilitate comparison to other atlasing efforts we report metrics from BigBrainSym, but make the re-registered segmentation available.

### MRI

#### Datasets

Three *in vivo* 7-Tesla MRI datasets with isotropic resolutions of 0.5mm, 0.7mm, and 1.0mm were analysed, each comprising 10 unique healthy adult participants. The 0.5mm “MICA-PNI” dataset was acquired in Montreal, Canada, and is publicly available^38^ (https://osf.io/mhq3f/). The 0.7mm and 1.0mm datasets were acquired at Maastricht University, The Netherlands, and were previously published^44,54^. All participants were healthy adults with no history of major neurological illness. Demographic details are provided in **Table 2**. These isotropic resolutions were chosen as 0.5mm represents the upper bound of whole-brain *in vivo* resolution presently achievable within reasonable scan times (<15 minutes); 0.7mm is achieved by recent, large public datasets^101^; and 1.0mm remains the most typical resolution of structural MRI, even among recent claustrum studies (**Fig. 1**). Note that classical sampling adequacy criteria such as the 5% voxel-to-ROI volume guideline are satisfied at all resolutions but prove misleading for thin structures^64^.

**Table 2.** Demographic details of three MRI datasets.

| Dataset resolution<br>(mm isotropic) | Scanner location | N | Sex (female) | Age (mean, SD) |
| --- | --- | --- | --- | --- |
| 0.5 | Montreal | 10 | 6 | 26.60 (4.60) |
| 0.7 | Maastricht | 10 | 4 | 28.60 (4.17) |
| 1.0 | Maastricht | 10 | 5 | 25.70 (2.94) |

#### Acquisition

The 0.5mm dataset was acquired on a Siemens 7-Tesla Magnetom Terra (Siemens Healthineers, Erlangen, Germany), using a 32Rx/8Tx head coil (NOVA Medical Inc., Wilmington, MA, United States). For this dataset, three runs were obtained at separate time points over an average span of 96.45 (±74.71) days. The 0.7mm and 1.0mm datasets were acquired in one run on a Siemens 7-Tesla Magnetom using a 32Rx/1Tx head coil (NOVA Medical Inc., Wilmington, MA, United States). From all datasets, we utilised whole-brain 3D-MP2RAGE uniform (UNI) images (T1-weighted)^102^, on which the claustrum appears hypointense. Acquisition details are provided in **Table 3**.

**Table 3.** Acquisition parameters of three MRI datasets.

| Dataset resolution<br>(mm isotropic) | TE (ms) | TR (ms) | Flip angle (°) | Tl<sub>1</sub>/ Tl<sub>2</sub> (ms) | Scan length<br>(m:s) | Acceleration<br>factor (PE) |
| --- | --- | --- | --- | --- | --- | --- |
| 0.5 | 2.44 | 5170 | 4/4 | 1000/3200 | 12:35 | 3 |
| 0.7 | 2.47 | 5030 | 5/3 | 900/2750 | 8:07 | 3 |
| 1.0 | 2.35 | 4500 | 5/3 | 900/2750 | 7:14 | 3 |

#### Pre-processing

All participants’ MP2RAGE UNI images were visually inspected for artifacts (e.g., ghosting, Gibbs ringing) and adequate subcortical contrast, and deemed suitable for inclusion. All images underwent background noise removal and bias-field correction using AFNI^103^ via in-house tools (https://github.com/srikash/3dMPRAGEise), and skull-stripping using SynthStrip^104^ via ‘mri_synthstrip’ in Freesurfer v7.4.1^105^. Despite sufficient signal-to-noise in the individual runs, we constructed an unbiased average template from the three 0.5mm runs to further improve effective signal and anatomical stability, using ANTs v2.4.4^106^, with six degrees of freedom and normalised mutual information as the cost function, though the claustrum was similarly identifiable in individual runs. The 0.5mm template (averaged across three runs) and the single-scan 0.7mm and 1mm datasets were used for all subsequent analyses.

#### Processing

To quantify differences in claustrum visibility across the three MRI datasets, we calculated the contrast-to-noise ratio (CNR), defined as the absolute difference in mean intensity between the segmented claustrum and its surrounding white matter, normalised by the standard deviation of the white matter signal^107^. In each dataset, approximately 60mm³ of white matter voxels were selected from the left hemisphere extreme and external capsulae in the coronal view using ITK-SNAP. We also estimated intracranial volume (ICV) using the recon-all pipeline in FreeSurfer^108^, for use as a covariate in sex-difference analyses.

#### Segmentation approach

The claustrum was manually segmented in native space by a single rater (SP). Segmentations were performed in ITK-SNAP using a one-voxel-sized brush, with simultaneous visualisation of all three orthogonal planes and the three-dimensional volume. Segmentation was based solely on visibility in the MP2RAGE UNI contrast, without direct comparison to the histological gold standard. A differential approach to segmentation was applied across claustral subregions, following a protocol first developed for 0.7mm isotropic MRI, with the dorsal claustrum segmented primarily in the axial view, the ventral claustrum in the coronal view, and the sagittal view used primarily to validate the posterior temporal claustrum^69^. In light of prominent partial voluming at claustral boundaries, a liberal approach was taken in which hypointense voxels were included if judged to be primarily claustral, i.e., containing discernible grey matter, even when directly abutting other grey matter structures.

A second rater (NC) conducted full quality control, including manual refinements. Claustrum segmentations were verified to avoid overlap with cortical grey matter, as defined by the subject-specific cortical ribbon (ribbon.mgz) generated by the FreeSurfer recon-all pipeline^108^, and with subcortical structures, specifically the putamen and amygdala, manually annotated at 0.3mm isotropic resolution on BigBrain transformed to ICBMsym space using an improved registration protocol^59^. Additionally, the second rater fully segmented the left hemisphere of one subject from each dataset in duplicate, achieving high inter-rater agreement (average DSC=0.93, see **Supplementary Fig. 1**).

We opted for manual segmentation after testing automated segmentation algorithms. Automation is highly desirable not only to reduce time and expertise demands, but also to curb annotation “style” that may limit cross-study comparability. Yet the same thin-sheet geometry and partial voluming that challenge humans also confound algorithms: widely used whole-brain parcellation algorithms either perform poorly (BrainSuite; Nighres), conflate the claustrum with adjacent structures (e.g., FreeSurfer SAMSEG), or omit it entirely (e.g., SPM, FSL, AFNI). Five recent bespoke algorithms have specifically targeted the claustrum, either alone or alongside a small number of other subcortical structures^52,78,85,89,109^. We applied these five algorithms to our three MRI datasets, but found that for each algorithm, in every dataset, automated segmentations were less consistent with manual segmentations than human raters were with each other, suggesting poor generalisation (**Supplementary Table 6)**.

#### Alignment and registration

##### Rigid alignment

Pre-processed MRI scans and native-space segmentations were rigidly aligned to the symmetric MNI ICBM152 nonlinear 2009b template^110^, using ANTsRegistration^106^, to correct for residual differences in head position that persist despite head stabilization and may bias morphometric measurements. All morphometric measurements (described below) were computed in this aligned space.

##### Non-linear registration

For voxel-wise comparisons required by spatial agreement metrics (described below), rigidly aligned images were further processed through a full affine and nonlinear registration pipeline. Affine registration (12 degrees of freedom) and symmetric diffeomorphic registration (SyN) to the symmetric MNI ICBM152 nonlinear 2009b template^110^ was performed using ANTs^111^, with a cross-correlation cost function. The resulting transformations were applied to segmentation labels using GenericLabel interpolation to preserve binary values. Registration accuracy was visually validated by overlaying each subject’s registered anatomy with subcortical structures defined by the Xiao atlas^59^. Warped claustrum segmentations were also inspected and found to be well-aligned, with occasional minor deviations (≤1 voxel) consistent with expected interpolation effects and the structural thinness of the claustrum. To maintain reproducibility, no manual corrections were applied to warped labels.

### Downsampled histology

To evaluate the effect of spatial resolution, the gold standard segmentation was downsampled to ‘MRI-like’ isotropic resolutions ranging from 0.4mm to 2.0mm (in 0.1mm increments) across several thresholds (0.2–0.8), using FSL’s ‘flirt’ with trilinear interpolation in three dimensions^112,113^. By comparing the downsampled gold standard to segmentations derived from acquired MRI, we effectively test whether spatial resolution alone accounts for observed differences. Substantial discrepancies would suggest that additional factors, such as contrast differences between histological staining and T1-weighted MR imaging, contribute to the difficulty of capturing the claustrum *in vivo*. See **Supplementary Fig. 4** for an example of downsampling effects.

### Morphometric measurements

#### Three dimensional metrics

We computed six three-dimensional metrics to characterise claustrum segmentations from the histological gold standard, rigidly aligned MRI datasets, and the downsampled gold standards. *Volume* was calculated as the number of labelled voxels multiplied by voxel resolution, reported in cubic millimetres. *Extents* were calculated as the maximal dimension along each orthogonal axis (x, y, z), in millimetres. *Roundness* (dimensionless) was computed as a ratio comparing the surface area of a sphere with the same Feret diameter as the segmentation’s mesh, where values near 1 indicate a spherical shape and lower values reflect increasingly elongated or irregular geometry. *Flatness* (dimensionless) was calculated as the square root of the ratio between the structure’s second-smallest and smallest eigenvalues, with larger values indicating more planar, sheet-like structures. To complement *Extents*, we computed the *Oriented Bounding Box (OBB)*, the minimal bounding box (x′, y′, z′) enclosing each claustrum irrespective of axis alignment, in millimetres. OBB was excluded from statistical comparisons to reduce the number of multiple comparisons. All metrics were computed over three-dimensional volumes using the ‘Label Map Statistics’ module in 3D Slicer (v5.6.2)^114^. We did not report absolute surface area, as it is ill-defined for structures without a closed surface representation, nor did we normalise metrics by intracranial volume (other than for the analysis of sex differences).

#### Two dimensional metrics

The claustrum’s thin mediolateral profile follows a curved, non-linear anatomical trajectory; as a result, three-dimensional metrics such as axis-aligned extents and oriented bounding boxes, which integrate across this curvature, can obscure the degree of thinness evident in individual two-dimensional slices. To better capture this property, we computed two two-dimensional (slice-wise) metrics in the coronal plane: ‘mean thickness, total voxel span’ as the distance between the minimum and maximum x-values of segmented voxels in each slice, irrespective of contiguity, and ‘mean thickness, contiguous voxels’ as the average width of all uninterrupted segments along the x-axis, capturing interruptions due to intervening white matter (see **Extended Data Fig. 3**). These two thickness measures diverge when claustrum segmentation becomes fragmented within individual coronal slices, with the ventral claustrum showing the greatest divergence, measured by the ratio between total span and contiguous thickness. Finally, for visualisation, we projected the three-dimensional gold standard segmentation along each orthogonal axis and summed voxel counts to generate flattened thickness maps. (This visualisation was not computed for MRI or the downsampled gold standards, as their spatial resolution is insufficient to distinguish anatomical thinness from voxel sampling effects due to partial voluming.)

### Objectives and statistical analysis

#### Objective 1. Characterising claustrum morphology across resolutions

Our first objective was to quantify how claustrum morphology varies across datasets that differ in spatial resolution and imaging modality: a high-resolution histological gold standard, its synthetically downsampled derivatives, and three rigidly-aligned *in vivo* MRI datasets. Analyses focused on eight morphometric metrics as defined above.

##### Analysis 1: Anatomy of the histological gold standard

First, we anatomically characterised the gold standard claustrum. Though based on a single brain, this high-resolution model preserves fine structural detail and serves as a reference for both the morphometric comparisons that follow and qualitative comparisons to prior anatomical reports.

##### Analysis 2: Resolution-dependent morphological degradation in downsampled gold standards

Next, we assessed how spatial resolution affects morphometric fidelity by downsampling the gold standard to a range of *in vivo* MRI-like resolutions. This simulated data, free of bias due to contrast or noise, define the theoretical maximum detail recoverable by MRI at each resolution. For each of the eight metrics (averaged across hemispheres), we fit a general linear model (GLM) with resolution and binarisation threshold as continuous fixed effects, including their interaction. Linear, quadratic, and cubic forms were tested, with the best-fitting model selected via likelihood ratio tests, Akaike Information Criterion (AIC), and Bayesian Information Criterion (BIC). Effect sizes were computed using adjusted R².

##### Analysis 3: The claustrum as captured by *in vivo* MRI

Finally, we assessed how morphometric estimates varied across the three *in vivo* MRI datasets. For each of the eight morphological metrics (averaged across hemispheres), we performed a one-way ANOVA with resolution as a fixed factor. Where significant effects were observed, pairwise comparisons were made using Tukey’s HSD *post hoc* tests, and effect sizes were reported using η².

To evaluate measurement stability within each dataset, we quantified intra-dataset variability using the coefficient of variation (CV), defined as the ratio of the standard deviation to the mean^115^. Differences in variability across datasets were assessed using Levene’s test, with Games–Howell *post hoc* comparisons for pairwise differences. We expected variability to increase at lower resolutions. Finally, to assess the impact of image quality, we tested whether CNR predicted segmentation variability by regressing CNR against each participant’s absolute deviation from the dataset mean^116^.

#### Objective 2. Evaluating MRI accuracy against histological and resolution-matched gold standards

Our second objective was to evaluate the degree of *in vivo* MRI capture by comparing segmentations to both the histological gold standard (anatomical “truth”) and its synthetically downsampled derivatives at matched resolutions (‘resolution ceiling’). In addition to comparing the same eight morphometric metrics defined above, we quantified spatial correspondence using four agreement metrics on MNI-aligned MRI segmentations. The Dice Similarity Coefficient (DSC)^117^ quantifies volumetric overlap, ranging from 0 (no overlap) to 1 (perfect agreement). Hausdorff Distance (HD)^118^ measures the greatest distance (mm) between the closest points on each segmentation boundary, capturing maximal misalignment, ranging from 0 (perfect alignment) to infinity. Given the claustrum’s high boundary-to-volume ratio (an upshot of its mediolateral thinness), and that standard spatial agreement metrics are known to penalise complex and thin structures^119^, we also computed dilated DSC (dDSC), which dilates and erodes each segmentation by one voxel prior to comparison, reducing sensitivity to minor boundary mismatches^92,120^, as well as balanced average HD (baHD), which normalises directional distances based on the number of ground truth points, mitigating bias introduced by differences in segmentation size^121^. Spatial agreement metrics were computed in MNI coordinates to ensure that spatial agreement reflects physical brain anatomy rather than voxel indices, enabling comparisons across datasets with different voxel resolutions.

##### Analysis 4: Morphometric and spatial agreement between MRI and histological gold standard

To assess how closely MRI segmentations approximated claustral morphology as revealed by histology, we compared the eight morphometric measurements from the three observed MRI datasets to the corresponding values derived from the histological gold standard. Deviation from the gold standard was described using percent differences, Cohen’s *d* effect sizes, and one-sample t-tests. Spatial correspondence was assessed using the four spatial agreement metrics.

##### Analysis 5: MRI performance relative to resolution-matched gold standards

To assess how closely MRI segmentations approached the theoretical limits imposed by their spatial resolution, we compared each MRI dataset to the corresponding downsampled gold standard binarised at a 50% threshold, which provides the theoretical ceiling. We chose a 50% threshold as we reasoned this is equivalent to a “majority-vote” rule, anchoring the ceiling in sampling physics rather than segmentation style. As in Analysis 4, we quantified deviation using percent differences and Cohen’s *d*, and evaluated spatial agreement using the same four metrics. Then, to quantify how much of the theoretically achievable DSC and HD agreement MRI attained at each resolution, we calculated ‘efficiency’ as the ratio of MRI performance to the theoretical ceiling: (MRI dataset vs. downsampled ÷ downsampled vs. gold standard) × 100% (note that HD efficiency required an inverted calculation as lower distances indicate better performance). We are not aware of efficiency analyses for other subcortical structures or thin structures—most validations report spatial agreement metrics with histology and/or *ex vivo* MRI without accounting for resolution-imposed ceilings^42,122^—so adopted what seemed like a fair albeit *post hoc* heuristic of ≥50-74% efficiency as adequate and ≥75-100% as high.

#### Exploratory analyses within MRI datasets

In addition to our primary objectives of describing claustrum anatomy and characterising the capacity to image it via MRI, we conducted three exploratory investigations using MRI data to address open questions in the literature. All analyses pooled data across the three MRI datasets (n=30). For all analyses, parametric tests were applied after verifying assumptions, and multiple comparisons were corrected using the Benjamini–Hochberg false discovery rate (FDR)^123^. Spatial agreement metrics were computed in Python (v3.11.4) using scipy^124^; all other statistical analyses were performed in R (v4.3.1).

##### Analysis 6: Inter-individual variability

To explore spatial variability in claustrum location across individuals, we created a probabilistic overlay from all MNI-aligned segmentations. Each dataset’s probability volume was resampled to the highest acquired resolution (0.5 mm isotropic) using trilinear interpolation, then averaged to produce a unified probability map. Voxel values represent the proportion of participants in whom the claustrum was present at each location, providing a spatial visualisation of inter-individual boundary consistency.

##### Analysis 7: Hemispheric asymmetry

To assess lateral differences in claustrum morphology, we analysed left and right claustra independently using paired-samples t-tests. We computed an asymmetry index (AI) for each participant as AI = (L – R)/(L + R)^125^, and used a GLM to test for dataset differences in AI, with ‘dataset’ included as a categorical covariate (0.5mm as reference).

##### Analysis 8: Sex differences

We analysed left and right claustra separately, assessing sex differences in each hemisphere using independent-samples t-tests. To account for known sex differences in total brain volume^91^, we then performed ANCOVA including intracranial volume (ICV) included as a covariate^126^.

## Left hemisphere

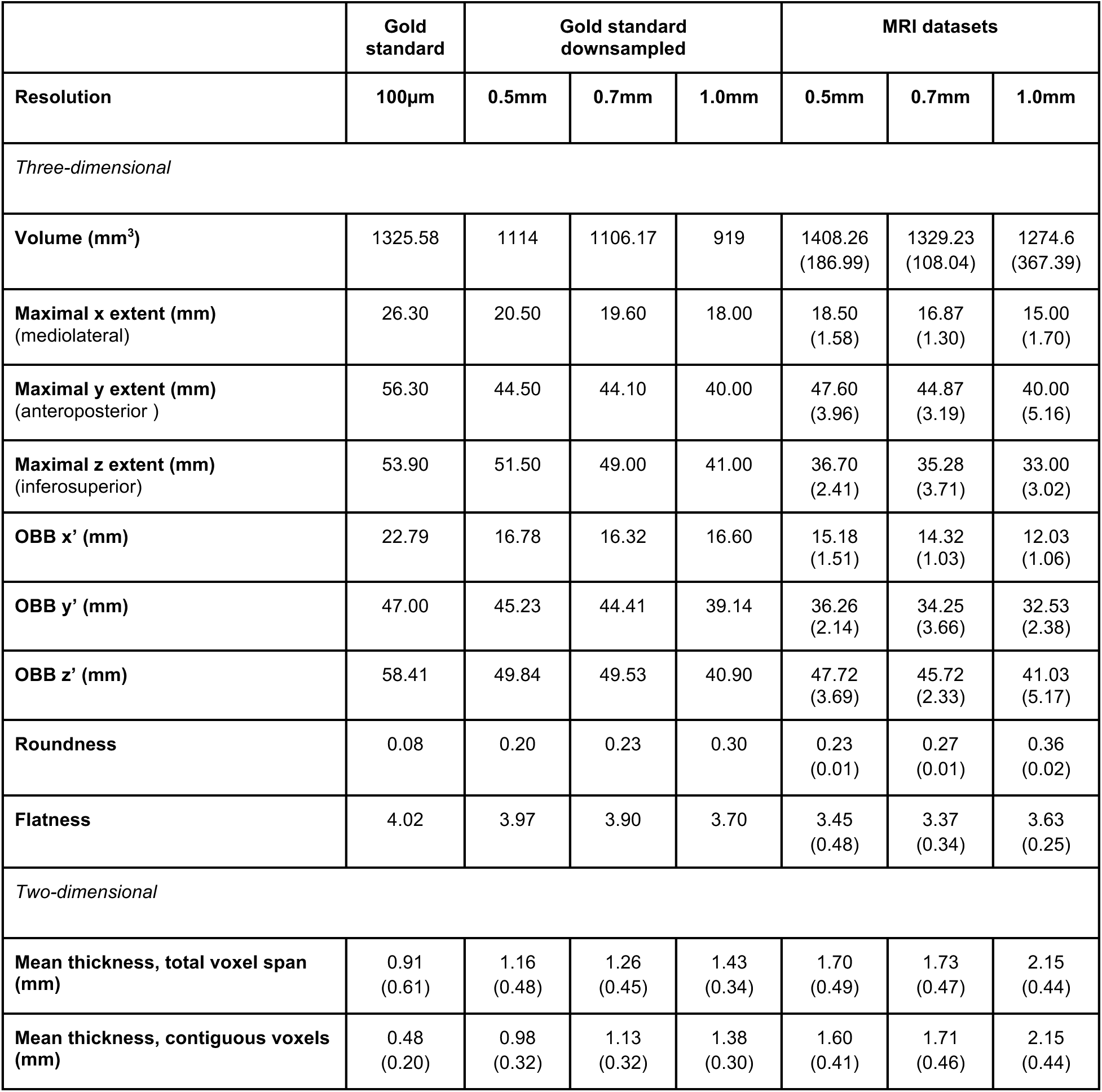

|  | Gold standard | Gold standard downsampled |  |  | MRI datasets |  |  |
| --- | --- | --- | --- | --- | --- | --- | --- |
| Resolution | 100µm | 0.5mm | 0.7mm | 1.0mm | 0.5mm | 0.7mm | 1.0mm |
| *Three-dimensional* |  |  |  |  |  |  |  |
| Volume (mm<sup>3</sup>) | 1325.58 | 1114 | 1106.17 | 919 | 1408.26<br>(186.99) | 1329.23<br>(108.04) | 1274.6<br>(367.39) |
| Maximal x extent (mm)<br>(mediolateral) | 26.30 | 20.50 | 19.60 | 18.00 | 18.50<br>(1.58) | 16.87<br>(1.30) | 15.00<br>(1.70) |
| Maximal y extent (mm)<br>(anteroposterior) | 56.30 | 44.50 | 44.10 | 40.00 | 47.60<br>(3.96) | 44.87<br>(3.19) | 40.00<br>(5.16) |
| Maximal z extent (mm)<br>(inferosuperior) | 53.90 | 51.50 | 49.00 | 41.00 | 36.70<br>(2.41) | 35.28<br>(3.71) | 33.00<br>(3.02) |
| OBB x' (mm) | 22.79 | 16.78 | 16.32 | 16.60 | 15.18<br>(1.51) | 14.32<br>(1.03) | 12.03<br>(1.06) |
| OBB y' (mm) | 47.00 | 45.23 | 44.41 | 39.14 | 36.26<br>(2.14) | 34.25<br>(3.66) | 32.53<br>(2.38) |
| OBB z' (mm) | 58.41 | 49.84 | 49.53 | 40.90 | 47.72<br>(3.69) | 45.72<br>(2.33) | 41.03<br>(5.17) |
| Roundness | 0.08 | 0.20 | 0.23 | 0.30 | 0.23<br>(0.01) | 0.27<br>(0.01) | 0.36<br>(0.02) |
| Flatness | 4.02 | 3.97 | 3.90 | 3.70 | 3.45<br>(0.48) | 3.37<br>(0.34) | 3.63<br>(0.25) |
| *Two-dimensional* |  |  |  |  |  |  |  |
| Mean thickness, total voxel span (mm) | 0.91<br>(0.61) | 1.16<br>(0.48) | 1.26<br>(0.45) | 1.43<br>(0.34) | 1.70<br>(0.49) | 1.73<br>(0.47) | 2.15<br>(0.44) |
| Mean thickness, contiguous voxels (mm) | 0.48<br>(0.20) | 0.98<br>(0.32) | 1.13<br>(0.32) | 1.38<br>(0.30) | 1.60<br>(0.41) | 1.71<br>(0.46) | 2.15<br>(0.44) |

## Right hemisphere

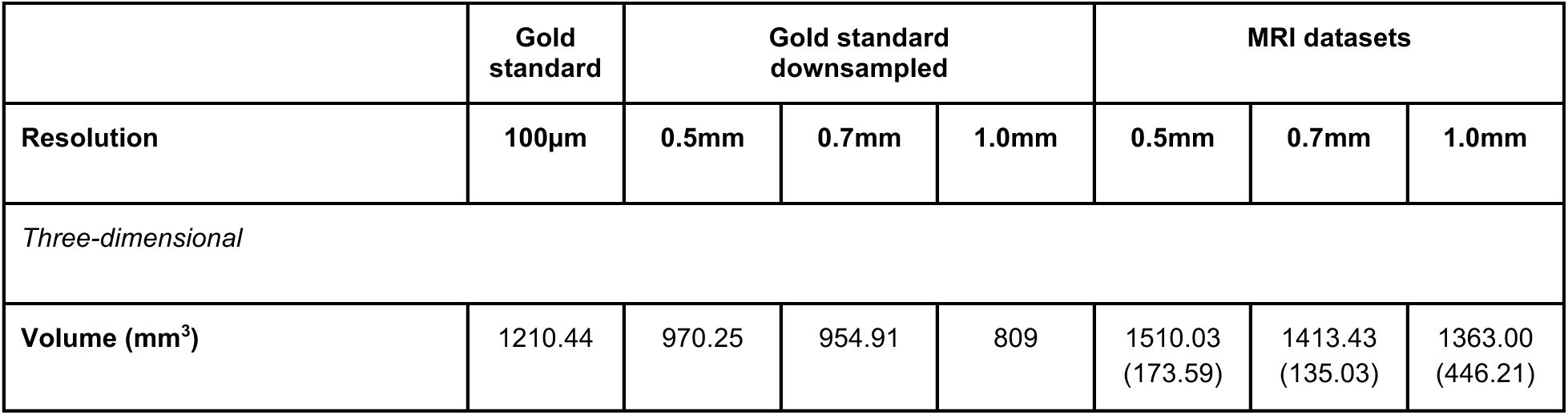

---

**Extended Data Table 1.**
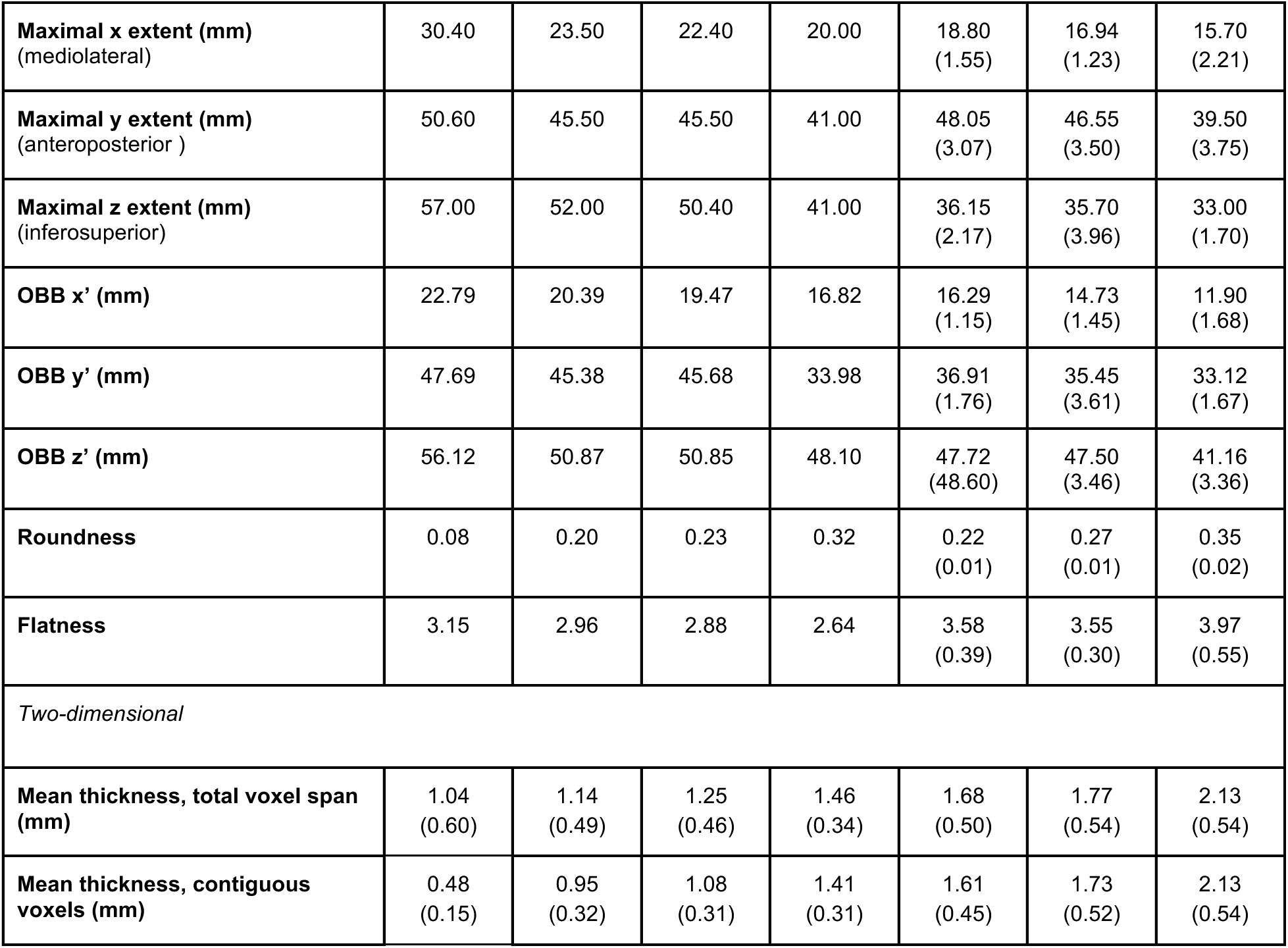
Hemisphere-specific morphometric measurements from gold standard, downsampled gold standards, and MRI datasets. Downsampled gold standards thresholded at 50%. For MRI datasets, values are averaged across participants, and standard deviations reflect inter-subject variability. For the gold standard and its downsampled versions (single brain), no standard deviation is reported, except for two-dimensional thickness metrics, where values are averaged across coronal slices and standard deviation reflects within-structure variation.

**Extended Data Table 2.** Statistical comparison of the MRI datasets on all morphometric measurements.

| | Corrected<br>p value | $\eta^2$ | Significant pairwise differences<br>(p value) |
| --- | --- | --- | --- |
| *Three-dimensional* |  |  |  |
| **Volume (mm<sup>3</sup>)** | 0.246 | 0.048 | *None* |
| **Maximal x extent (mm)**<br>(mediolateral) | <0.001 | 0.428 | 0.5mm vs 0.7mm = 0.003<br>0.5mm vs 1.0mm < 0.001<br>0.7mm vs 1.0mm = 0.009 |
| **Maximal y extent (mm)**<br>(anteroposterior) | <0.001 | 0.464 | 0.5mm vs 1.0mm < 0.001<br>0.7mm vs 1.0mm < 0.001 |
| **Maximal z extent (mm)**<br>(inferosuperior) | 0.002 | 0.211 | 0.5mm vs 1.0mm = 0.001<br>0.7mm vs 1.0mm = 0.022 |
| **Roundness** | <0.001 | 0.945 | 0.5mm vs 0.7mm < 0.001<br>0.5mm vs 1.0mm < 0.001<br>0.7mm vs 1.0mm < 0.001 |
| **Flatness** | 0.025 | 0.126 | 0.7mm vs 1.0mm = 0.027 |
| *Two-dimensional* |  |  |  |
| **Mean thickness, total voxel span (mm)** | <0.001 | 0.691 | 0.5mm vs 1.0mm = < 0.001<br>0.7mm vs 1.0mm = < 0.001 |
| **Mean thickness, contiguous voxels (mm)** | <0.001 | 0.758 | 0.5mm vs 0.7mm = 0.026<br>0.5mm vs 1.0mm < 0.001<br>0.7mm vs 1.0mm < 0.001 |

**Extended Data Table 3.** Differences between gold standard and MRI datasets. Positive percent differences indicate MRI values exceed gold standard values. All p-values were corrected using false discovery rate (FDR) correction across 24 comparisons.

|  | Gold standard vs MRI datasets |  |  |  |  |  |  |  |  |
| --- | --- | --- | --- | --- | --- | --- | --- | --- | --- |
|  | Percent difference |  |  | Cohen's *d* |  |  | Corrected p value |  |  |
| Resolution (mm) | 0.5 | 0.7 | 1.0 | 0.5 | 0.7 | 1.0 | 0.5 | 0.7 | 1.0 |
| *Three dimensional* |  |  |  |  |  |  |  |  |  |
| **Volume (mm<sup>3</sup>)** | +15.30 | +8.36 | +4.26 | 1.00 | 0.75 | 0.12 | <0.001 | 0.004 | 0.608 |
| **Maximal x extent (mm)**<br>(mediolateral) | -33.72 | -40.03 | -45.58 | -3.17 | -4.53 | -4.59 | <0.001 | <0.001 | <0.001 |
| **Maximal y extent (mm)**<br>(anteroposterior ) | -10.25 | -14.15 | -25.45 | -1.20 | -1.55 | -2.70 | <0.001 | <0.001 | <0.001 |
| **Maximal z extent (mm)**<br>(inferosuperior) | -34.29 | -35.97 | -40.45 | -7.96 | -5.13 | -8.16 | <0.001 | <0.001 | <0.001 |
| **Roundness** | +186.05 | +242.40 | +344.79 | 13.95 | 17.09 | 16.17 | <0.001 | <0.001 | <0.001 |
| **Flatness** | -0.57 | -1.97 | +7.07 | -0.12 | -0.22 | 0.46 | 0.608 | 0.360 | 0.063 |
| *Two-dimensional* |  |  |  |  |  |  |  |  |  |
| **Mean thickness, total voxel span (mm)** | +74.30 | +80.03 | +120.81 | 5.84 | 5.58 | 6.36 | <0.001 | <0.001 | <0.001 |
| **Mean thickness, contiguous voxels (mm)** | +234.3 | +257.41 | +343.74 | 10.78 | 9.75 | 10.55 | <0.001 | <0.001 | <0.001 |

**Extended Data Table 4.** Agreement between gold standard, downsampled gold standard, and MRI datasets.

|  | Gold standard vs.<br>downsampled gold standard |  |  | Gold standard vs. MRI |  |  | Downsampled gold standard<br>vs. MRI |  |  |
| --- | --- | --- | --- | --- | --- | --- | --- | --- | --- |
|  | 0.5 | 0.7 | 1.0 | 0.5 | 0.7 | 1.0 | 0.5 | 0.7 | 1.0 |
| **DSC** | 0.82<br>(0.02) | 0.81<br>(0.02) | 0.71<br>(0.02) | 0.39<br>(0.02) | 0.40<br>(0.03) | 0.37<br>(0.03) | 0.41<br>(0.03) | 0.44<br>(0.03) | 0.41<br>(0.04) |
| **HD (mm)** | 6.28<br>(0.23) | 6.80<br>(2.34) | 13.28<br>(0.67) | 9.49<br>(2.35) | 12.12<br>(2.99) | 13.05<br>(2.23) | 8.29<br>(1.55) | 9.98<br>(3.10) | 10.65<br>(1.78) |
| **dDSC** | 0.82<br>(0.02) | 0.83<br>(0.03) | 0.70<br>(0.03) | 0.59<br>(0.02) | 0.63<br>(0.02) | 0.61<br>(0.03) | 0.60<br>(0.02) | 0.66<br>(0.03) | 0.64<br>(0.05) |
| **baHD (mm)** | 0.16<br>(0.02) | 0.22<br>(0.05) | 0.69<br>(0.08) | 0.71<br>(0.07) | 0.83<br>(0.16) | 1.18<br>(0.13) | 0.71<br>(0.09) | 0.75<br>(0.15) | 1.15<br>(0.29) |
| **Jaccard (IoU)** | 0.69<br>(0.02) | 0.68<br>(0.02) | 0.55<br>(0.02) | 0.24<br>(0.02) | 0.26<br>(0.02) | 0.23<br>(0.02) | 0.26<br>(0.02) | 0.28<br>(0.02) | 0.26<br>(0.03) |

**Extended Data Table 5.** Differences between resolution-matched downsampled gold standards (50% threshold) and MRI. Positive percent differences indicate MRI values exceed downsampled gold standard values. All p-values were corrected using false discovery rate (FDR) correction across 24 comparisons.

|  | Downsampled gold standards vs MRI datasets |  |  |  |  |  |  |  |  |
| --- | --- | --- | --- | --- | --- | --- | --- | --- | --- |
|  | Percent difference |  |  | Cohen's *d* |  |  | Corrected p value |  |  |
| Resolution (mm) | 0.5 | 0.7 | 1.0 | 0.5 | 0.7 | 1.0 | 0.5 | 0.7 | 1.0 |
| *Three-dimensional* |  |  |  |  |  |  |  |  |  |
| **Volume (mm<sup>3</sup>)** | +40.66 | +33.76 | +53.34 | 2.32 | 2.71 | 1.16 | <0.001 | <0.001 | <0.001 |
| **Maximal x extent (mm)**<br>(mediolateral) | -14.64 | -19.11 | -19.00 | -2.14 | -3.21 | -1.89 | <0.001 | <0.001 | <0.001 |
| **Maximal y extent (mm)**<br>(anteroposterior ) | +6.29 | +2.03 | -1.83 | 0.84 | 0.28 | -0.17 | 0.002 | 0.260 | 0.504 |
| **Maximal z extent (mm)**<br>(inferosuperior) | -29.62 | -28.59 | -19.51 | -6.99 | -3.89 | -3.44 | <0.001 | <0.001 | <0.001 |
| **Roundness** | +13.83 | +18.19 | +15.27 | 2.78 | 3.77 | 2.89 | <0.001 | <0.001 | <0.001 |
| **Flatness** | +3.53 | +4.47 | +22.42 | 0.11 | 0.20 | 1.34 | 0.731 | 0.624 | <0.001 |
| *Two-dimensional* |  |  |  |  |  |  |  |  |  |
| **Mean thickness, total voxel span (mm)** | +47.27 | +39.56 | +48.21 | 4.65 | 3.94 | 4.49 | <0.001 | <0.001 | <0.001 |
| **Mean thickness, contiguous voxels (mm)** | +66.61 | +55.98 | +53.49 | 6.30 | 4.97 | 4.86 | <0.001 | <0.001 | <0.001 |

**Extended Data Fig. 1.**
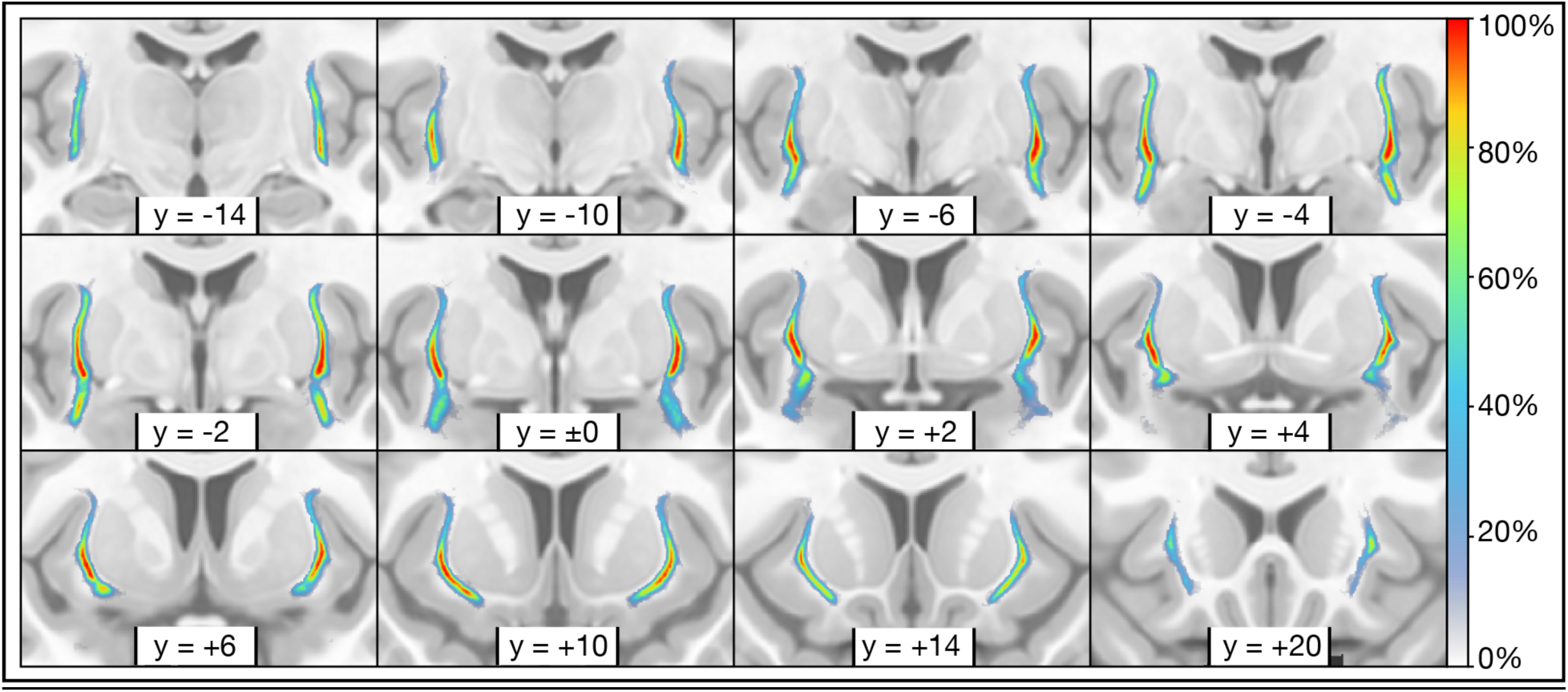
Probabilistic overlay of MRI datasets. Coronal slices show voxel-wise overlap of MNI-aligned claustrum segmentations from all three MRI datasets (n=30), resampled to 0.5 mm isotropic resolution. Voxel intensity reflects the proportion of participants with claustrum present at each location (0–100%). A consistent central core spans the anteroposterior extent (Y = −14 to +20mm shown), with highest agreement in the dorsal midsection. Variability increases toward the periphery, particularly ventrally and anteriorly, reflecting reduced thickness and greater boundary ambiguity.

**Extended Data Fig. 2.**
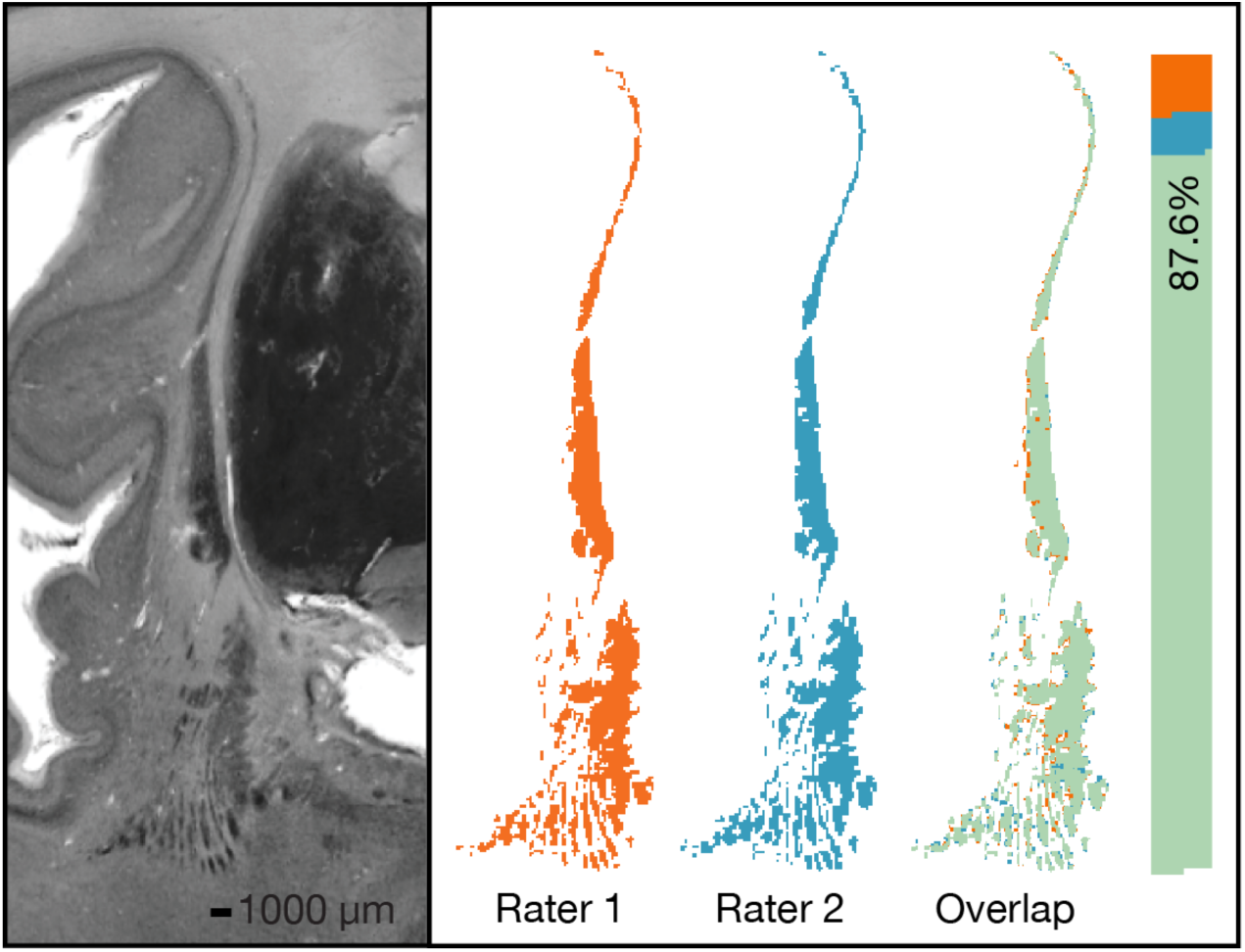
Inter-rater agreement of gold standard claustrum segmentation. Inter-rater agreement was assessed via duplicate segmentation of three randomly selected coronal slices in the right hemisphere, spaced ≥75 slices apart and containing >100 voxels in both segmentations. Rater 2 segmented de novo without access to Rater 1’s work. Dice Similarity Coefficient (DSC) ranged from 0.87 to 0.93, indicating high inter-rater agreement. Best-case agreement shown for a single slice (BigBrain y=1350, DSC=0.93). Left: BigBrain histology. Right: Segmentations by Rater 1 (SP, orange, 6100 voxels) and Rater 2 (NC, blue, 5960 voxels), with overlap (green) and unique voxels in respective colors. Bar shows agreement (87.6%) and disagreement (12.4%). Disagreements primarily occurred along edges and in ventral “puddles.“

**Extended Data Figure 3.**
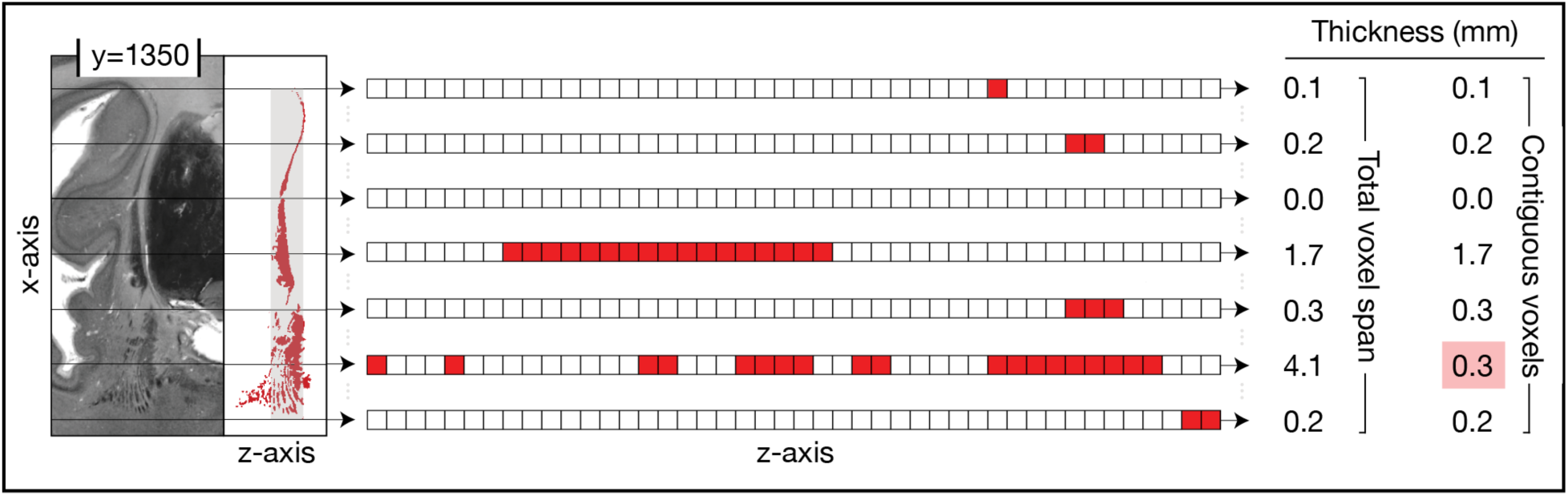
Slice-wise calculations to capture claustral thinness. Two slice-wise metrics were computed across the anteroposterior extent to quantify claustral thinness. The illustration shows how ‘mean thickness, total voxel span’, and ‘mean thickness, contiguous voxels’ were calculated for an example coronal slice (y=1350) of the right hemisphere of the gold standard, near the claustrum’s midpoint. Left: the histological image and corresponding segmentation (red) illustrate variation in claustral thickness in two dimensions along the x-axis. This variability is further compounded in three dimensions, as the claustrum follows a curved trajectory from anterior to posterior. Middle: seven equidistant positions along the x-axis (of 455 total) are highlighted. Right: the table shows counts for both metrics, and highlights (pink) differences in the ventral claustrum. Mean thickness of contiguous voxels, which adjusts for white matter interruptions, is particularly relevant for MRI where partial voluming may cause ventral “puddles” to fall below detection thresholds or appear artefactually thickened. In the slice shown, the mean total voxel span was 2.46 mm, while the mean thickness of contiguous voxels was 1.16 mm (ratio=2.12).

**Supplementary Table 1.** MNI coordinates of claustrum centre of mass. Centre of mass coordinates (x, y, z) for left and right claustra across the gold standard and MRI datasets, in MNI space (mm). MRI-derived centres closely approximate the gold standard, with most falling within several voxel’s distance.

|  |  | Left hemisphere coordinates (mm) |  |  | Right hemisphere coordinates (mm) |  |  |
| --- | --- | --- | --- | --- | --- | --- | --- |
| Dataset type | Dataset resolution (mm isotropic) | x | y | z |  | y | z |
| BigBrain | 0.1 | -32.32 | +0.90 | -5.37 | 31.68 | 1.09 | -6.01 |
| MRI | 0.5 | -32.45 (0.53) | 1.43 (0.88) | -3.03 (1.81) | 32.09 (0.19) | 2.38 (0.78) | -3.57 (1.93) |
| MRI | 0.7 | -32.56 (0.28) | 0.83 (0.84) | -2.67 (1.82) | 32.14 (0.24) | 1.90 (1.10) | -4.04 (1.86) |
| MRI | 1.0 | -33.26 (0.62) | -0.24 (1.24) | -2.35 (1.78) | 31.99 (0.46) | 1.49 (1.30) | -3.69 (1.82) |

**Supplementary Table 2.** Coefficient of variation (CV) of morphometric measurements in MRI datasets. Variability across participants within each MRI dataset. All metrics showed low variability (CV < 0.15) except volume at 1.0 mm (CV = 0.31), indicating reduced measurement stability at lower resolution.

| Resolution (mm) | 0.5mm | 0.7mm | 1.0mm |
| --- | --- | --- | --- |
| *Three-dimensional* |  |  |  |
| **Volume (mm<sup>3</sup>)** | 0.12 | 0.08 | 0.31 |
| **Maximal x extent (mm)**<br>(mediolateral) | 0.06 | 0.06 | 0.10 |
| **Maximal y extent (mm)**<br>(anteroposterior ) | 0.06 | 0.10 | 0.05 |
| **Maximal z extent (mm)**<br>(inferosuperior) | 0.06 | 0.06 | 0.11 |
| **Roundness** | 0.04 | 0.04 | 0.06 |
| **Flatness** | 0.10 | 0.08 | 0.08 |
| *Two-dimensional* |  |  |  |
| **Mean thickness, total voxel span (mm)** | 0.07 | 0.07 | 0.07 |
| **Mean thickness, contiguous voxels (mm)** | 0.06 | 0.07 | 0.07 |

**Supplementary Table 3.** Ratio between total voxel span and contiguous thickness for each dataset, computed across the full claustrum and within the middle third of the anteroposterior axis. The gold standard shows large discrepancies, whereas MRI ratios approach 1.00, reflecting resolution-driven loss of anatomical detail.

|  | Gold standard | Gold standard downsampled |  |  | MRI datasets |  |  |
| --- | --- | --- | --- | --- | --- | --- | --- |
| Resolution | 100μm | 0.5mm | 0.7mm | 1.0mm | 0.5mm | 0.7mm | 1.0mm |
| Span-to-contiguous thickness ratio: full extent | 1.85 (0.79) | 1.16 (0.20) | 1.11 (0.17) | 1.04 (0.09) | 1.05 (0.10) | 1.01 (0.05) | 1.00 (0.01) |
| Span-to-contiguous thickness ratio: middle third | 2.76 (0.58) | 1.35 (0.21) | 1.28 (0.20) | 1.08 (0.13) | 1.08 (0.10) | 1.02 (0.04) | 1.00 (0.01) |
| Maximum ratio: full extent | 4.29 | 1.86 | 1.69 | 1.43 | 2.40 | 1.68 | 1.20 |

**Supplementary Table 4.** MRI performance efficiency relative to theoretical limits. Dice similarity coefficient (DSC) and Hausdorff distance (HD) efficiency for each MRI dataset, defined as the proportion of achievable volumetric overlap or boundary precision recovered relative to the theoretical ceiling (downsampled vs. gold standard). See also **Fig. 8**.

|  | Downsampled gold standard vs. MRI |  |  |
| --- | --- | --- | --- |
| Resolution (mm) | 0.5 | 0.7 | 1.0 |
| DSC efficiency (%) | 50.00 | 54.32 | 57.75 |
| HD efficiency (%) | 75.75 | 68.14 | 124.69 |

**Supplementary Table 5.** Claustrum morphometrics for super-high resolution *ex vivo* MRI (100µm; single brain)^70^, independently segmented by Coates & Zaretskaya^51^ and Mauri and colleagues^52^. Gold standard values are included for comparison. Values reflect the average across hemispheres; bracketed values reflect inter-hemispheric differences, not standard deviations.

|  |  | Edlow MGH dataset (2019) |  |
| --- | --- | --- | --- |
|  | Gold standard | Coates & Zaretskaya (2024) | Mauri and colleagues (2025) |
| **Resolution** | **100µm** | **100µm** | **100µm** |
| *Three-dimensional* |  |  |  |
| **Volume (mm<sup>3</sup>)** | 1268.01<br>(81.42) | 1905.32<br>(239.12) | 1453.46<br>(21.78) |
| **Maximal x extent (mm)**<br>(mediolateral) | 28.35<br>(2.90) | 25.65<br>(4.31) | 19.95<br>(0.21) |
| **Maximal y extent (mm)**<br>(anteroposterior ) | 53.45<br>(4.03) | 53.70<br>(7.50) | 45.10<br>(1.41) |
| **Maximal z extent (mm)** (inferosuperior) | 55.45<br>(2.19) | 51.35<br>(1.20) | 40.85<br>(3.32) |
| **OBB x'** | 24.41<br>(2.30) | 20.59<br>(4.78) | 15.33<br>(1.05) |
| **OBB y'** | 47.35<br>(0.49) | 53.72<br>(1.64) | 43.18<br>(3.40) |
| **OBB z'** | 57.26<br>(1.62) | 53.39<br>(5.01) | 46.45<br>(1.00) |
| **Roundness** | 0.08<br>(0.00) | 0.16<br>(0.05) | 0.17<br>(0.02) |
| **Flatness** | 3.58<br>(0.62) | 4.40<br>(0.79) | 4.48<br>(0.56) |
| *Two-dimensional* |  |  |  |
| **Mean thickness, total voxel span (mm)** | 0.97<br>(0.60) | 1.16<br>(0.64) | 1.14<br>(0.56) |
| **Mean thickness, contiguous voxels (mm)** | 0.48<br>(0.17) | 1.07<br>(0.52) | 1.09<br>(0.50) |
| **Span-to-contiguous thickness ratio: full extent** | 1.85<br>(0.79) | 1.08<br>(0.21) | 1.04<br>(0.08) |
| **Span-to-contiguous thickness ratio: middle third** | 2.76<br>(0.58) | 1.20<br>(0.31) | 1.10<br>(0.10) |

**Supplementary Table 6.** Testing of automated claustrum segmentation algorithms. Manual claustrum segmentations of the 0.5mm dataset compared to five automated algorithms for adult brains^52,78,85,89,109^. Agreement was assessed using Dice Similarity Coefficient (DSC), Hausdorff Distance (HD), dilated DSC (dDSC), and balanced average HD (baHD). Values are mean (SD) across n=10 participants. Brun and Mauri’s algorithms were developed for 7-Tesla; others 3T. All algorithms except Mauri’s were trained on lower resolution data than that to which we applied them here (Brun=0.6mm, Berman=0.7mm, Albishri=0.7mm, and Li=1.0mm, all isotropic voxels). Note that Berman’s method is designed for dorsal claustrum only.

| Algorithm | DSC | HD | dDSC | baHD |
| --- | --- | --- | --- | --- |
| Albishri (2020) | 0.21 (0.17) | 88.12 (14.94) | 0.26 (0.19) | 48.24 (8.36) |
| Berman (2022) | 0.42 (0.10) | 30.67 (4.45) | 0.46 (0.09) | 21.59 (3.22) |
| Brun (2021) | 0.69 (0.02) | 16.48 (3.96) | 0.74 (0.03) | 11.75 (2.44) |
| Li (2022) | 0.47 (0.10) | 80.31 (21.22) | 0.50 (0.10) | 39.79 (7.26) |
| Mauri (2025) | 0.62 (0.02) | 13.59 (1.70) | 0.72 (0.02) | 11.29 (1.39) |

**Supplementary Fig. 1.**
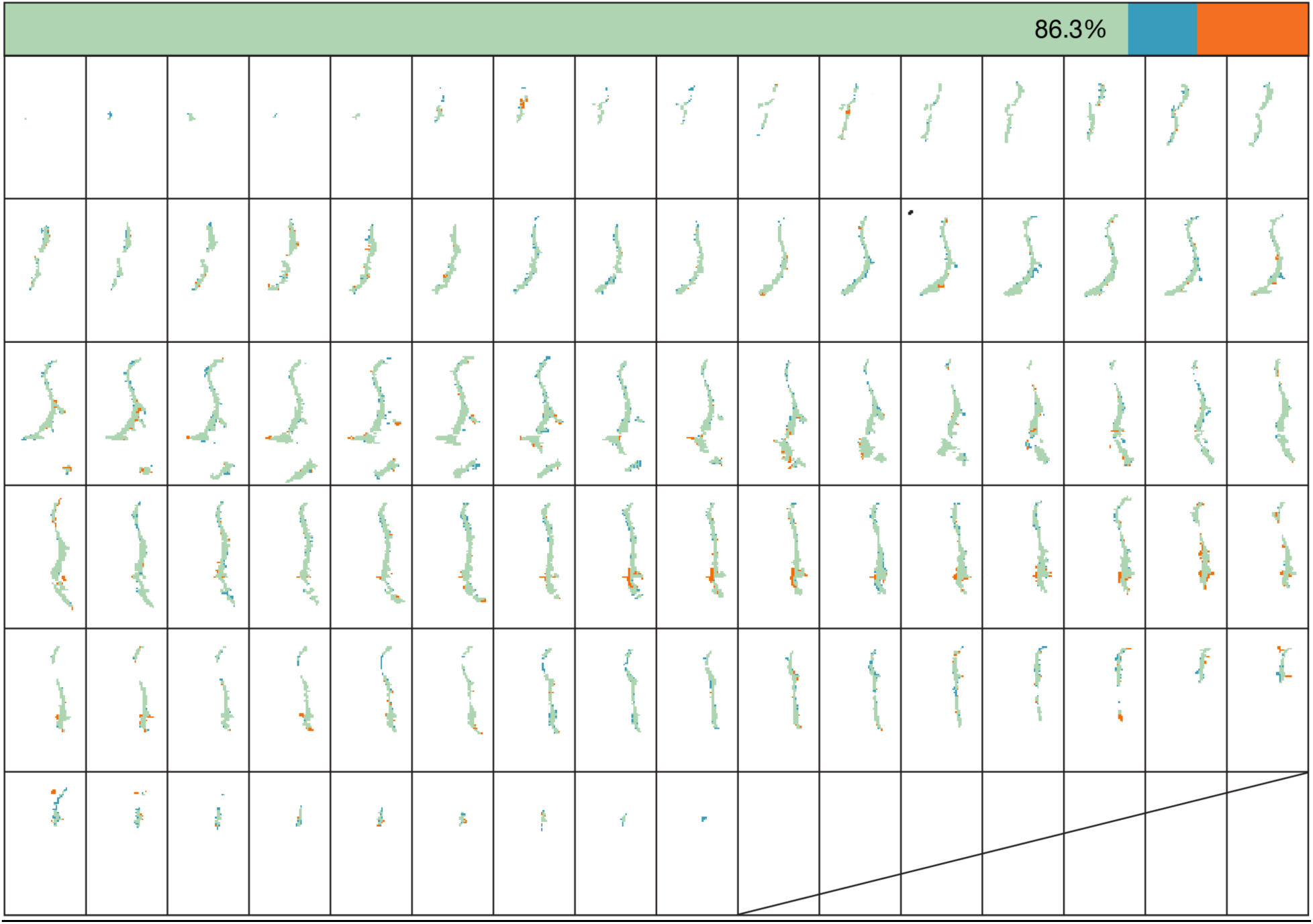
Inter-rater agreement of MRI segmentation. The left hemisphere from the participant with the most average volume in each dataset was segmented independently by two raters. Agreement assessed using Dice Similarity Coefficient (DSC) showed high structural overlap at all resolutions: DSC = 0.926 at 0.5mm, 0.941 at 0.7mm, and 0.934 at 1.0mm isotropic. Coronal slices (anterior-to-posterior) show left hemisphere segmentations from both raters for the 0.5mm dataset participant with lowest agreement (DSC = 0.926). Rater 1 (SP, orange), Rater 2 (NC, blue), and overlap (green); voxels segmented by only one rater shown in their respective color. Horizontal bar shows proportions of agreement (86.3%) and disagreement (13.7%). Consistent with gold standard segmentation (**Extended Data Fig. 2**), disagreements occurred primarily along edges and in the ventral claustrum.

**Supplementary Fig. 2.**
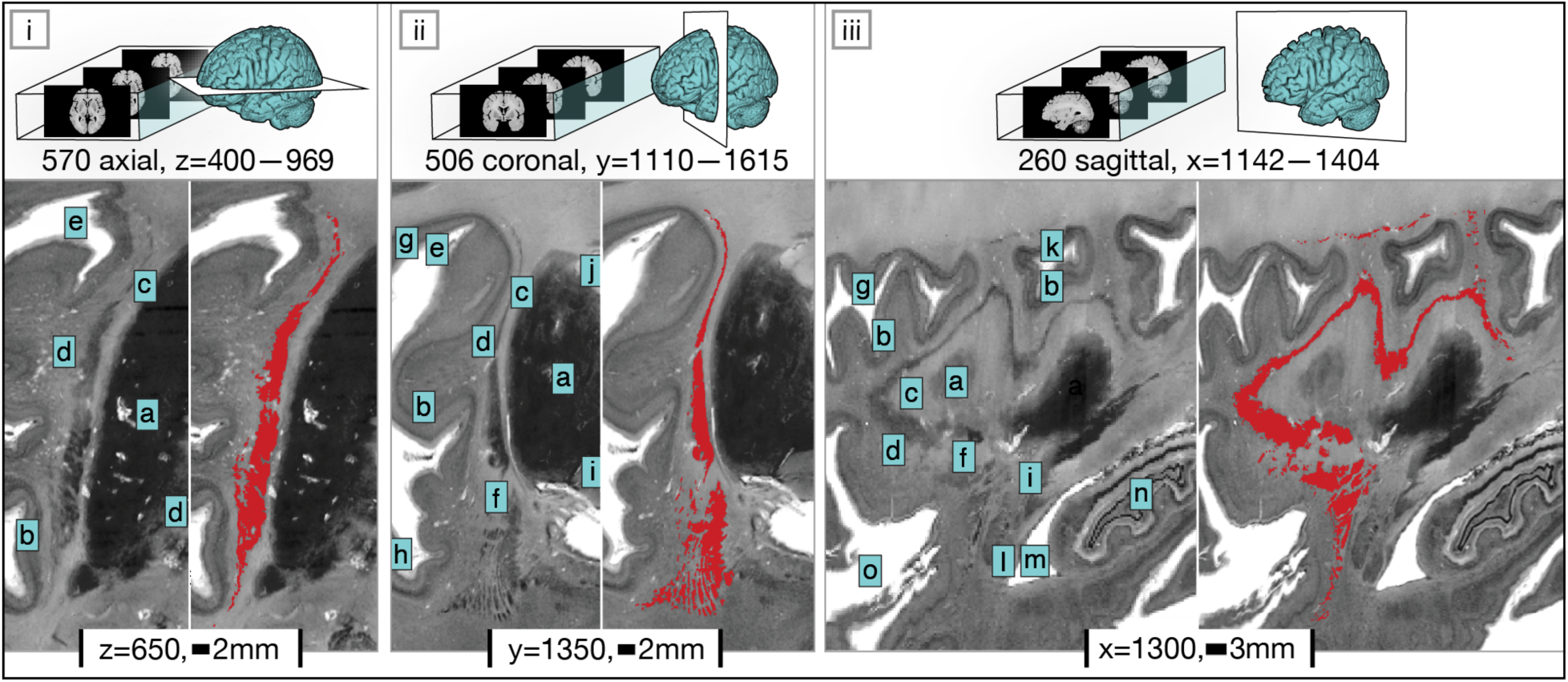
**BigBrain-derived gold standard claustrum model**. The right hemisphere claustrum delineation is shown, with vertical panels corresponding to approximately midpoint (i) axial, (ii) coronal, and (iii) sagittal views. The top row indicates the number of slices with a claustrum label, and visualises the location of the slice shown below (BigBrain coordinate given). The bottom row displays cropped BigBrain (left) alongside the corresponding claustrum label in red (right), highlighting the extraordinary detail achieved via slice-wise manual segmentation with a one-voxel brush. Letters mark nearby structures and spaces: (a) putamen, (b) insular cortex, (c) external capsule, (d) extreme capsule, (e) circular sulcus, (f) uncinate fascicle, (g) frontal operculum, (h) planum temporale, (i) anterior commissure, (j) internal capsule, (k) parietal operculum, (l) lateral amygdaloid nucleus, (m) lateral ventricle, (n) hippocampus, (o) lateral sulcus.

**Supplementary Fig. 3.**
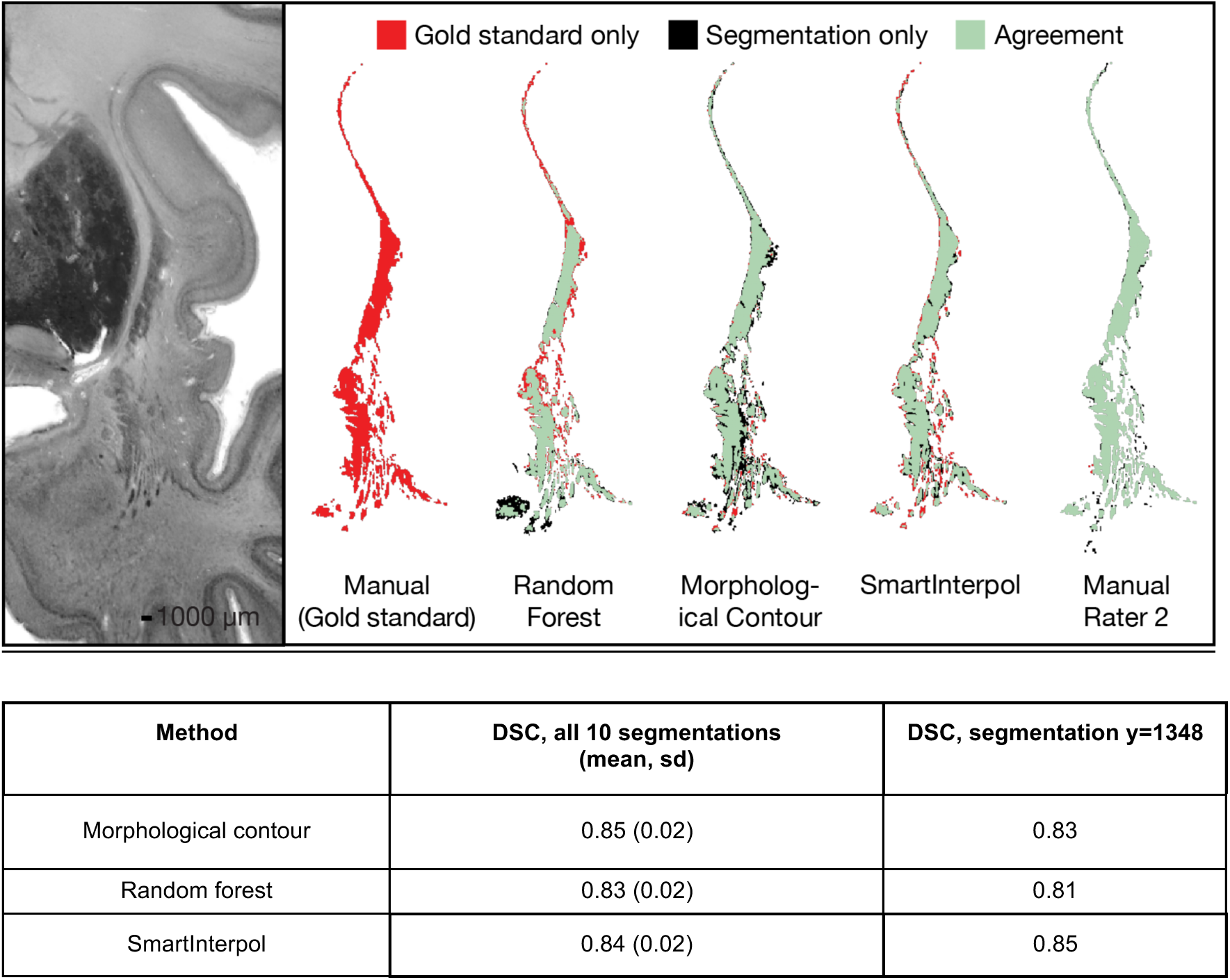
Testing 3 automated segmentation algorithms on histology. We tested three sparse interpolation algorithms with default parameters to evaluate their potential for reducing manual segmentation workload: morphological contour ^98^, random forest ^99^, and SmartInterpol ^100^ (using the product rule segmentation, which combines label fusion and deep learning). In a test region encompassing the ventral claustrum as it extends into the temporal lobe (28 consecutive coronal slices, BigBrain coordinates y=1335-1365), we manually segmented all slices but provided only every third slice (including the first and last) to each algorithm. On the task of segmenting interleaved 10 segmentations, all three methods produced good agreement with manual segmentation (see Table, below). In contrast, two human raters achieved excellent agreement (DSC=0.97) on a test slice (y=1348) on which all algorithms showed just good agreement. Lower algorithmic performance may stem from the claustrum’s highly undulating morphology between slices, violating the algorithms’ assumptions of high inter-slice correlation. Certainly, all methods would likely show improved results with tuning, but for challenging regions like the ventral claustrum, we judged that manual segmentation was essential and remains best practice. The higher human inter-rater agreement observed here (compared to that reported in **Extended Data Fig. 2**) may be because Rater 2 was provided with the same sparse input as the algorithms; in the earlier comparison, segmentation was performed *de novo*.

**Supplementary Fig. 4.**
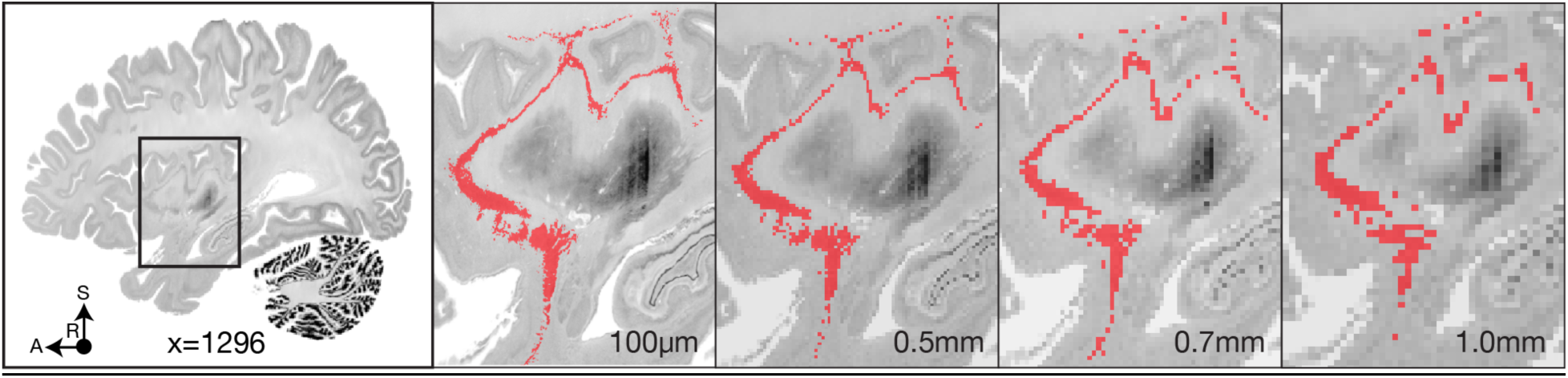
Downsampling analysis. Left: Inset shows a sagittal view of the BigBrain dataset (slice x=1296) in the right hemisphere, with a box indicating the zoomed region shown in subsequent panels. Right: The first panel displays the gold standard claustrum segmentation at 100μm resolution (red), followed by the same segmentation after downsampling to resolutions matched to the three acquired MRI datasets, thresholded at 50%. The comparison illustrates how spatial resolution affects anatomical detail: while gross shape and topology are preserved at submillimetric levels, finer features are progressively lost at lower resolutions.

## Supplementary Note 1. Suggestions for reporting

Our results motivate reporting standards to make claustrum findings interpretable and comparable across studies:

1. Report nominal voxel size and effective resolution at the capsule-claustrum boundary in the mediolateral direction, and interpret both against the histological gold standard’s mean contiguous mediolateral thickness (∼0.56mm).
2. Specify the slice plane and its obliquity relative to AC-PC and to an insula-aligned oblique-coronal plane parallel to the extreme and external capsules.
3. Report claustrum-to-capsular CNR.
4. Describe the segmentation protocol (manual or semi-automatic), any initialisation (for example, warping the gold standard for localisation guidance), inter- and intra-rater reliability, and any *post hoc* topology corrections.
5. Segment in native space; for group analyses, describe the non-linear registration and any local refinement near the claustrum, as thin structures are highly sensitive to warp error and topology breaks.
6. State explicitly which features visible in the gold standard were not detectable with MRI; if some participants were differentially affected (e.g., with ventral “drop out”), consider exclusion criteria based on per-subject claustral capture, though this risks non-random missingness.
7. Report morphometrics beyond volume; we recommend the eight two-dimensional and three-dimensional metrics used here.

## Supplementary Note 2. Creation of cross-modal, probabilistic claustrum atlas

To facilitate claustrum identification and mitigate the limiation that the gold standard derives from a single histological specimen, we created a probabilistic atlas that combines the gold standard and 0.5mm *in vivo* MRI dataset described in the present paper, together with the super-high resolution *ex vivo* and *in vivo* MRI segmentations released by Mauri and colleagues^52^:

| Dataset resolution (mm isotropic) | Dataset | N | Segmentation provided by | Modality | Acquisition | Total Weight |
| --- | --- | --- | --- | --- | --- | --- |
| 0.10 | BigBrain<sup>53</sup> | 1 | This paper | Histology | *Ex vivo* | 20% |
| 0.10 | Edlow MGH brain<sup>70</sup> | 1 | Mauri (2025)<sup>52</sup> | 7T MRI | *Ex vivo* | 20% |
| 0.25 | Lüsebrink brain<sup>83</sup> | 1 | Mauri (2025)<sup>52</sup> | 7T MRI | *In vivo* | 20% |
| 0.50 | 0.5mm dataset<sup>54</sup> | 10 | This paper | 7T MRI | *In vivo* | 40% |

To generate the atlas in standard space, all claustrum segmentations were aligned to the MNI ICBM152 nonlinear 2009b template^110^ at 0.5mm isotropic resolution. For the two *ex vivo* datasets, publicly available MNI-aligned versions (BigBrain: https://osf.io/xkqb3/overview; Edlow MGH brain: https://datadryad.org/dataset/doi:10.5061/dryad.119f80q) were used as registration references as they provided superior subcortical alignment. Both *in vivo* datasets were registered directly to the MNI template, following the procedures outlined in the ‘Non-linear registration’ subsection of the Methods. Differential weights were applied such that higher-resolution datasets exerted greater influence on the final voxelwise probabilities.

The probabilistic atlas is shown overlaid on the MNI template^110^, with voxel intensities representing the weighted likelihood of claustral tissue at each location:

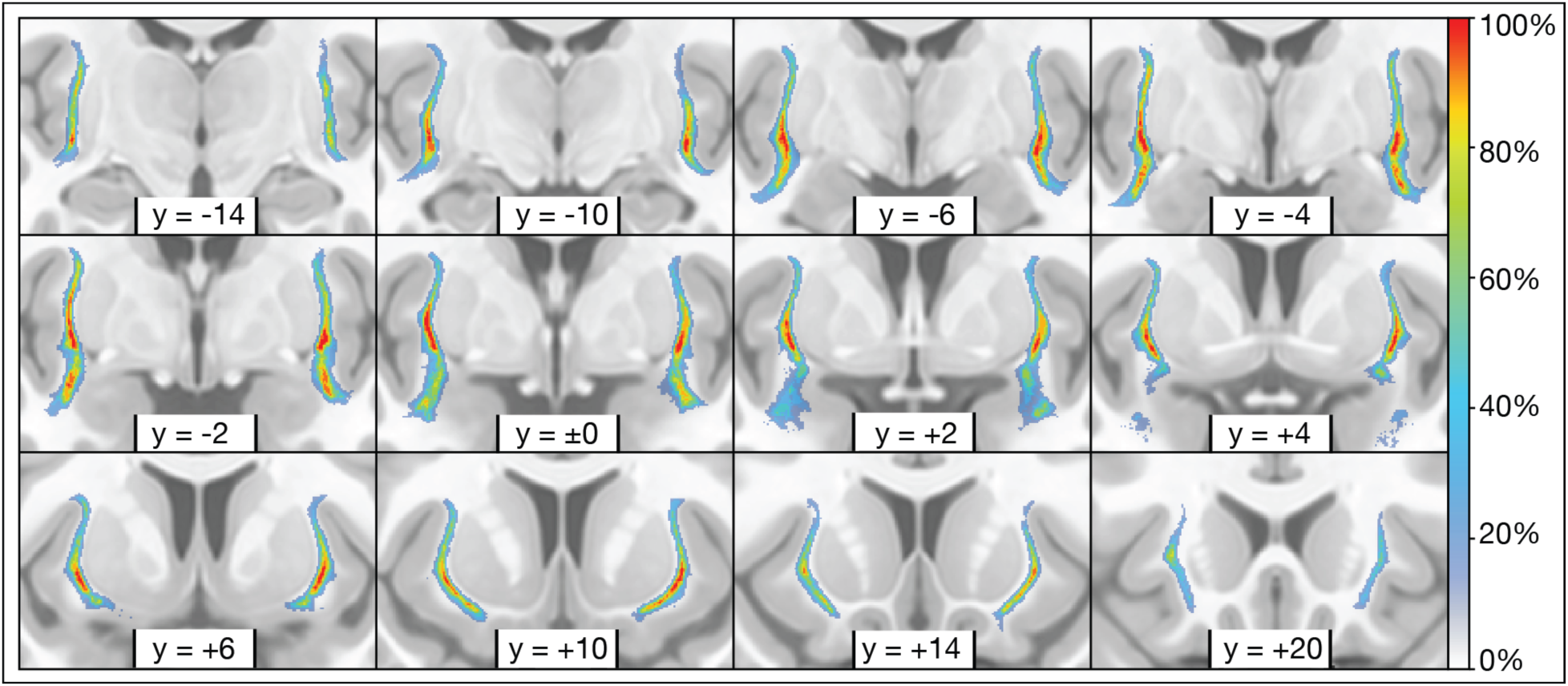

## Supplementary Note 3. 8-step quality control process for gold standard segmentation

I. Raters simultaneously observed labeling in all three planes (axial, coronal, and sagittal) alongside real-time three-dimensional volumetric reconstruction in ITK-SNAP.
II. Following Kang’s protocol developed for high-resolution MRI^69^, raters preferentially labeled aspects of the claustrum in specific views: dorsal regions in the axial view, ventral regions in the coronal view, and the sagittal view was consulted primarily for quality control.
III. Approximately every 25mm along the anteroposterior extent, and as needed to resolve ambiguity, raters cross-referenced their label with the BigBrain dataset at 20µm in-plane resolution^68^.
IV. The BigBrain dataset at 1µm in-plane resolution^68^ was also cross-referenced to ensure that the claustrum label did not overlap with existing labels of nearby structures, including the putamen, amygdala, and insular cortex.
V. Upon completion of the initial segmentation, the alternate rater performed a slice-by-slice quality control review of the opposite hemisphere, correcting clear errors and resolving notable discrepancies through discussion.
VI. Within the claustrum label, voxels with intensity values more than two standard deviations below the average labeled voxel contrast were flagged. These voxels were manually reviewed by the original rater and removed as necessary to limit the erroneous inclusion of white matter and blood vessels.
VII. The claustrum label was inflated by three voxels, and voxels with intensity values greater than the average labeled voxel contrast were flagged. These voxels were manually reviewed by the original rater and included as necessary to ensure consistent gray matter inclusion along edges.
VIII. Three randomly selected coronal slices in the right hemisphere were fully and independently labeled by the alternate rater, allowing for the measurement of inter-rater agreement.

## Notes

https://github.com/navonacalarco/claustrum

### Competing Interest Statement

The authors have declared no competing interest.

